# Acute malaria dysregulates specialized lymph node macrophages to suppress vaccine-elicited protection against Ebola virus

**DOI:** 10.1101/2025.09.10.675493

**Authors:** Jonah Elliff, Lindsey Grady, Kyle L. O’Donnell, Caitlin Messingham, Kai J. Rogers, Jobaida Akther, Andrew Thurman, Rahul Vijay, Alejandro Pezzulo, Troy Randall, Andrea Marzi, Noah S. Butler, Wendy Maury

## Abstract

The filovirus, Ebola virus (EBOV), causes outbreaks of Ebola virus disease (EVD) throughout equatorial Africa. ERVEBO® (rVSV/EBOV) is a replication-competent, recombinant vesicular stomatitis virus (rVSV)-vectored vaccine licensed to control EVD outbreaks. EVD outbreaks occur in regions endemic for *Plasmodium*-caused malaria. *Plasmodium* infections persist due in part to the parasite’s ability to evade sterilizing immunity which also dampens immune responses to heterologous vaccines. Acute murine *Plasmodium* infection at the time of rVSV/EBOV vaccination reduced vaccine-mediated protection against mouse-adapted EBOV (ma-EBOV) challenge. Decreased protection was associated with a *Plasmodium-*induced interferon gamma (IFN-γ)-mediated decrease of rVSV/EBOV replication in lymph node (LN) macrophages, resulting in reduced primary anti-EBOV glycoprotein antibody responses. Higher doses of rVSV/EBOV partially overcame the antibody deficits and elicited protective responses. Evidence of the negative impact of *Plasmodium* on the efficacy of low dose rVSV/EBOV vaccine protocols supports the use of high antigen loads in effective management of EVD outbreaks.

**IMPORTANCE:** We show that blood-stage murine *Plasmodium* infections negatively impacts the primary antibody response elicited by low dose rVSV/EBOV vaccination and results in reduced protection against a lethal dose of ma-EBOV. This defect occurs within the draining lymph node due to the elevation of IFN-γ elicted in *Py-*infected mice. The *Py-*imposed decrease in vaccine-mediated protection can be overcome with higher doses of rVSV/EBOV. While the strong protection conferred by rVSV/EBOV and significant side effects known to be associated with this vaccine have led to the suggestion that the vaccine dosage be reduced^19^, our studies provide a rationale for maintaining a higher dose.

## INTRODUCTION

Ebola virus (EBOV) belongs to the virus family *Filoviridae*; a group of enveloped negative-sense RNA viruses that can be highly pathogenic and cause episodic outbreaks of severe hemorrhagic fever^1,2^. EBOV causes the most filovirus-associated human disease in outbreaks of Ebola virus disease (EVD) with observed mortality rates that range as high as 90% depending on the outbreak context^2,3^. There are two vaccines currently recommended by the World Health Organization (WHO) and available for use in EVD outbreak responses. Both vaccines elicit antibodies that target the EBOV glycoprotein (EBOV GP), which are known to be protective against EVD^4–8^. ERVEBO^®^ (rVSV/EBOV) is a vaccine approved by the US Food and Drug Administration (FDA), composed of replication-competent, recombinant vesicular stomatitis virus (rVSV) that encodes the EBOV GP in place of the native VSV GP^9,10^. The other recommended vaccine is a two-dose regimen of replication-incompetent adenovirus serotype 26 encoding EBOV GP (Ad26.ZEBOV; Zabdeno) and vaccinia Ankara-vectored EBOV GP (MVA-BN-Filo; Mvabea) that is approved by the European Medicines Agency (EMA)^11,12^. Vaccination with replication-competent rVSV/EBOV has advantages over other vaccine platforms as it can quickly induce robust and protective EBOV GP-specific antibody responses with a single dose, a feature that is critical for management of EVD outbreaks^13–15^. Testing of rVSV/EBOV via vaccination of close contacts during the largest outbreak of EVD in West Africa in 2014-2016 demonstrated 100% efficacy in preventing EVD^15^, leading to rapid pre-qualification of the vaccine and subsequent use in outbreak responses. Findings from a robust observational study of rVSV/EBOV efficacy during the 2018 outbreak in the Democratic Republic of the Congo (DRC) were compatible with the initial studies, showing that rVSV/EBOV was 84% protective against EVD if administered 10 or more days before symptom onset^16^.

The currently licensed dose of rVSV/EBOV is 2×10^7^ plaque-forming units (PFU)^15,17^, however, one prior study demonstrated that as little as 10 PFU rVSV/EBOV delivered 4 weeks prior to challenge to naïve non-human primates (NHP) were fully protective against a lethal dose of EBOV^18^, further underlining the high efficacy of rVSV/EBOV. These findings have led to the suggestion that use of lower doses of rVSV/EBOV could remain efficacious while circumventing the significant adverse events associated with vaccine administration^19^. However, factors such as parasitic co-infections may render low doses of rVSV/EBOV insufficient for the optimal management of EVD outbreaks.

EVD outbreaks occur in geographical locations that have the highest burdens of malaria in the world^20^. Serologic studies showed nearly universal exposure of adults to *P. falciparum* in endemic areas^21^. The large burden of *Plasmodium* is highlighted in one cross-sectional investigation in Sierra Leone that demonstrated approximately 40% of the population were actively parasitemic at the time of the study^22^. Host immune responses to *Plasmodium* are complicated by the varied developmental stages of the parasite^23^. The bite of an infected mosquito inoculates the skin with a motile form of the parasite, which then circulates to the liver. Following clinically silent cell division and differentiation in liver hepatocytes, parasites are released into circulation to initiate a cycle of invasion and asexual replication in red blood cells (RBCs). Acute blood-stage infection elicits clinical symptoms and systemic inflammation, including the production of TNF-α and IFN-γ^24–26^. Clearance of blood-stage parasites and resolution of clinical disease is dependent on antibodies^27,28^. However, antibody responses develop slowly and incompletely in infected individuals, only resulting in clinical immunity after years of repeated infections^29,30^. This inefficient acquisition of immunity can be explained by diverse mechanisms including allelic variation^31^ and dysregulated inflammation^32^; these mechanisms converge to constrain the development of protective parasite-specific antibodies.

The impact of dysregulated inflammation is not limited to immunity targeting the parasite, as concurrent infection with *Plasmodium* can dampen humoral immunity to co-infecting pathogens and numerous heterologous immunizations^33–36^. One study has examined the impact of malaria on rVSV/EBOV vaccine responses, finding that adult healthcare workers with asymptomatic malaria at the time of rVSV/EBOV vaccination had a modest decrease in neutralizing antibody responses at 3-, 6-, and 12-months following vaccination^37^. Yet all study participants had protective levels of antibodies, suggesting that the currently utilized dose is sufficient to mediate protection in *Plasmodium* infected recipients. However, given the impact of *Plasmodium* on efficacy of other vaccines and the high incidence of malaria in regions at risk for EVD outbreaks, we proposed that an acute, blood-stage *Plasmodium* infection would impact the protective efficacy of low dose rVSV/EBOV.

To test this, we utilized mouse models of blood-stage *Plasmodium* infection, intramuscular rVSV/EBOV vaccination, and mouse-adapted EBOV (ma-EBOV) and demonstrate that acute *Plasmodium* infection decreases the protective efficacy of low dose rVSV/EBOV against a challenge with ma-EBOV. Suboptimal responses were linked to a *Plasmodium-*elicited, interferon gamma (IFN-γ)-dependent decrease of rVSV/EBOV replication in a population of lymph node (LN) resident CD169^+^ macrophages. Loss of CD169^+^ macrophage-dependent virus replication resulted in lower viral antigen load and decreased generation of EBOV GP-specific germinal center (GC) responses, thereby reducing the magnitude of protective anti-EBOV antibody responses. High dose vaccination with rVSV/EBOV could overcome the *Plasmodium-* associated deficit to provide protection against lethal ma-EBOV challenge in *Plasmodium-* infected vaccine recipients. These studies highlight that vaccine strategies utilizing low doses of rVSV/EBOV may have reduced efficacy in acutely *Plasmodium* infected recipients and suggest that administration of high antigen loads can compensate for malaria-associated defects and support robust protection against disease and death caused by EBOV.

## RESULTS

### An acute, blood-stage *Plasmodium* infection decreases rVSV/EBOV-mediated protection against ma-EBOV challenge

To assess the impact of acute, blood-stage malaria on the protective efficacy of low dose rVSV/EBOV, C57BL/6 mice were intravenously (i.v.) administered RBCs (iRBCs) infected with the rodent parasite *Plasmodium yoelii,* 17XNL (*Py*). *Py* causes an acute, non-lethal infection in mice and models key aspects of *P. falciparum* in humans^38,39^. A control cohort was injected with PBS (non-infected). At day 6 following *Py* infection, mice were intramuscularly (i.m.) injected with 10^3^ PFU of rVSV/EBOV (low dose), 2.5×10^5^ PFU of rVSV/EBOV (high dose), or left non-vaccinated. 10^3^ PFU of rVSV/EBOV is a 20-fold lower dose than the licensed body weight-adjusted equivalent delivered to humans, while 2.5×10^5^ PFU exceeds the licensed equivalent dose by 12-fold and models a high dose vaccination. Cohorts of *Py-*infected and non-infected mice were challenged 45 days post-vaccination (dpv) with 10 focus forming units (FFU) of ma-EBOV **(Fig. 1A)**. Both groups of non-vaccinated mice, regardless of *Py* infection status, succumbed to ma-EBOV challenge **(Fig. 1B,C; indicated by open red squares or open black circles).** Vaccination with 10^3^ PFU rVSV/EBOV protected 100% of the non-infected mice against ma-EBOV challenge and significantly reduced titers of infectious virus in whole blood, liver and spleen at 4 days post-infection (dpi), as well as serum levels of soluble GP (sGP) **(Fig. 1B-D** and **S1; indicated by solid black circles)**. In contrast, only 70% of *Py-*infected 10^3^ PFU recipients were protected and all animals in this cohort experienced weight loss, indicating reduced protection **(Fig. 1B,C; indicated by solid red squares)**. Further, these mice had high ma-EBOV titers in multiple tissues and high levels of serum sGP at 4 dpi that were statistically indistinguishable from the non-vaccinated cohorts **(Fig. 1D** and **S1; indicated by solid red squares)**. These data support that an acute, blood-stage *Plasmodium* decreases protection elicited by low dose rVSV/EBOV in a mouse model.

**Figure 1:**
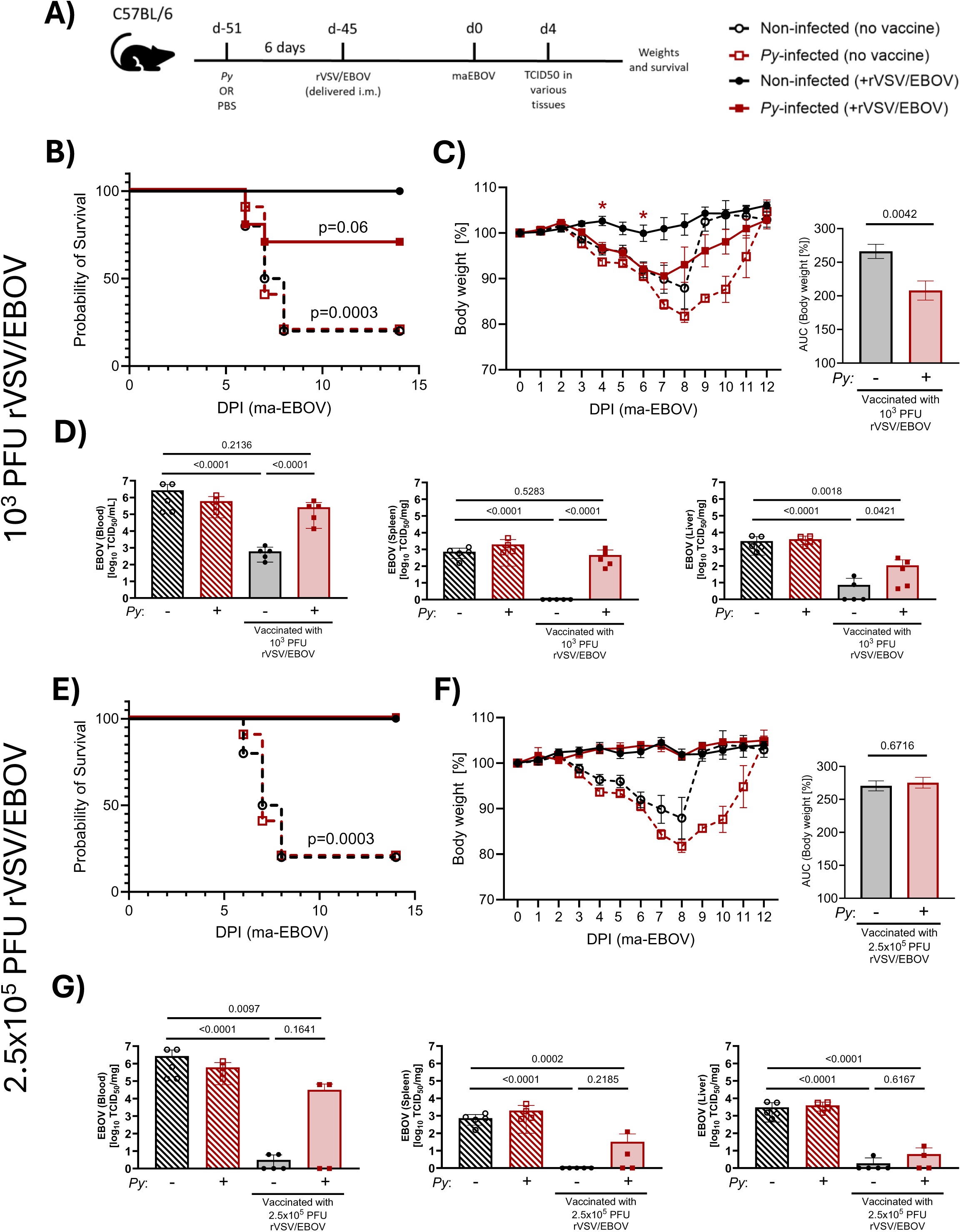
An acute, blood-stage *Plasmodium* infection decreases rVSV/EBOV-mediated protection against ma-EBOV challenge. (A) Female C57BL/6 mice were intravenously (i.v.) injected with PBS (non-infected; black circles) or infected with 1×10^6^ *Py*-infected red blood cells [iRBCs] (*Py*-infected; red squares). 6 days later, mice were vaccinated via intramuscular (i.m.) injection with 10^3^ PFU, 2.5×10^5^ PFU rVSV/EBOV, or left unvaccinated. Mice were challenged 45-dpv with 10 FFU mouse-adapted Ebola virus (ma-EBOV). Tissues were taken at 4-dpi (49-dpv) for ma-EBOV TCID_50_ and remaining mice were assessed for clinical signs and survival daily. (B, C) Survival curves and daily weights are shown (n=10/group) for 10^3^ PFU rVSV/EBOV recipients. Daily weights were evaluated by AUC for vaccine recipients. (D) Day 4 ma-EBOV TCID_50_ in whole blood, spleen and liver of 10^3^ PFU rVSV/EBOV recipients. (E, F) Survival curves and daily weights are shown (n=10/group) in 2.5×10^5^ PFU rVSV/EBOV recipients. Daily weights were evaluated by AUC for vaccine recipients. (G) Day 4 ma-EBOV TCID_50_ in whole blood, spleen and liver of 2.5×10^5^ PFU rVSV/EBOV recipients. Data for non-infected or *Py-*infected non-vaccinated controls are shared between A-C and D-E. ma-EBOV TCID_50_ are expressed as mean ± SD. p<0.05 considered significant. Data in B and E were analyzed by log-rank Mantel-Cox test. Data in C and F were analyzed using two-way ANOVA with Tukey’s multiple comparisons test, *p<0.05, and AUC was analyzed via students t-test and is represented as mean ± SEM. Data in D and G were analyzed using two-way ANOVA with Tukey’s multiple comparisons test. All data generated from one independent experiment. p<0.05 considered significant. Also see Figure S1.

Notably, all mice vaccinated with 2.5×10^5^ PFU rVSV/EBOV survived ma-EBOV challenge and had no observed weight loss, regardless of prior *Py* infection **(Fig. 1E,F)**. Infectious titers and serum sGP were also significantly decreased in *Py-*infected recipients of this dose compared to non-vaccinated cohorts **(Fig. 1G** and **S1),** although whole blood ma-EBOV titers in two of the four *Py-*infected high dose recipients were elevated **(Fig. 1G)**. Together, these data indicate that higher antigen loads can overcome the *Py-*imposed decrease rVSV/EBOV-mediated protection.

Collectively, these data demonstrate that an acute, blood-stage *Py* infection decreases protection mediated by low dose rVSV/EBOV vaccination in mice, while a high dose vaccination remains protective and reduces systemic ma-EBOV titers.

### Vaccine-elicited antibodies are decreased in *Plasmodium-*infected rVSV/EBOV recipients

Protection conferred by rVSV/EBOV is dependent on the production of anti-EBOV GP antibodies^4,5,8^. To determine whether the *Py-*imposed reduction in protection is a result of decreased antibody responses, mice with or without *Py* infection were vaccinated i.m. with 10^3^ PFU or 2.5×10^5^ PFU rVSV/EBOV. Serum was obtained at 21- and 42-dpv and anti-EBOV GP IgG was quantified **(Fig. 2A)**. Non-*Plasmodium* infected recipients of 10^3^ PFU rVSV/EBOV produced ∼100 µg/ml of anti-EBOV GP IgG, a circulating antibody concentration that we show here and in a previous study to be protective against ma-EBOV challenge^6^. In *Py*-infected recipients of this dose, serum anti-EBOV GP IgG was reduced by 12-fold at 21-dpv and 47-fold at 42-dpv **(Fig. 2B)**. We also observed a significant reduction in 21-dpv serum anti-EBOV GP IgG in BALB/c mice vaccinated with this dose **(Fig. S2A)**, demonstrating that *Py* elicits this deficit across different mouse strains. Vaccine administration of 2.5×10^5^ PFU into naïve C57BL/6 mice generated ∼300 µg/ml anti-EBOV GP IgG at 21 and 42-dpv **(Fig 2B)**. However, prior *Py* infection caused a ∼6-fold reduction in anti-EBOV IgG responses to this dose **(Fig. 2B),** a more modest *Py*-elicited deficit than that observed at the lower vaccine dose. Administration of an even higher dose of vaccine, 6.25×10^7^ PFU, still did not overcome the *Py*-induced antibody deficit as a ∼4-fold decrease was observed relative to non-infected vaccine recipients **(Fig S2B)**. Together, these data support that *Py* infection results in production of fewer rVSV/EBOV-elicited antibodies. Increasing the rVSV/EBOV dose can partially overcome these deficits, consistent with our observation that the high dose still confers protection when administered to *Py-*infected mice **(Fig 1).**

**Figure 2:**
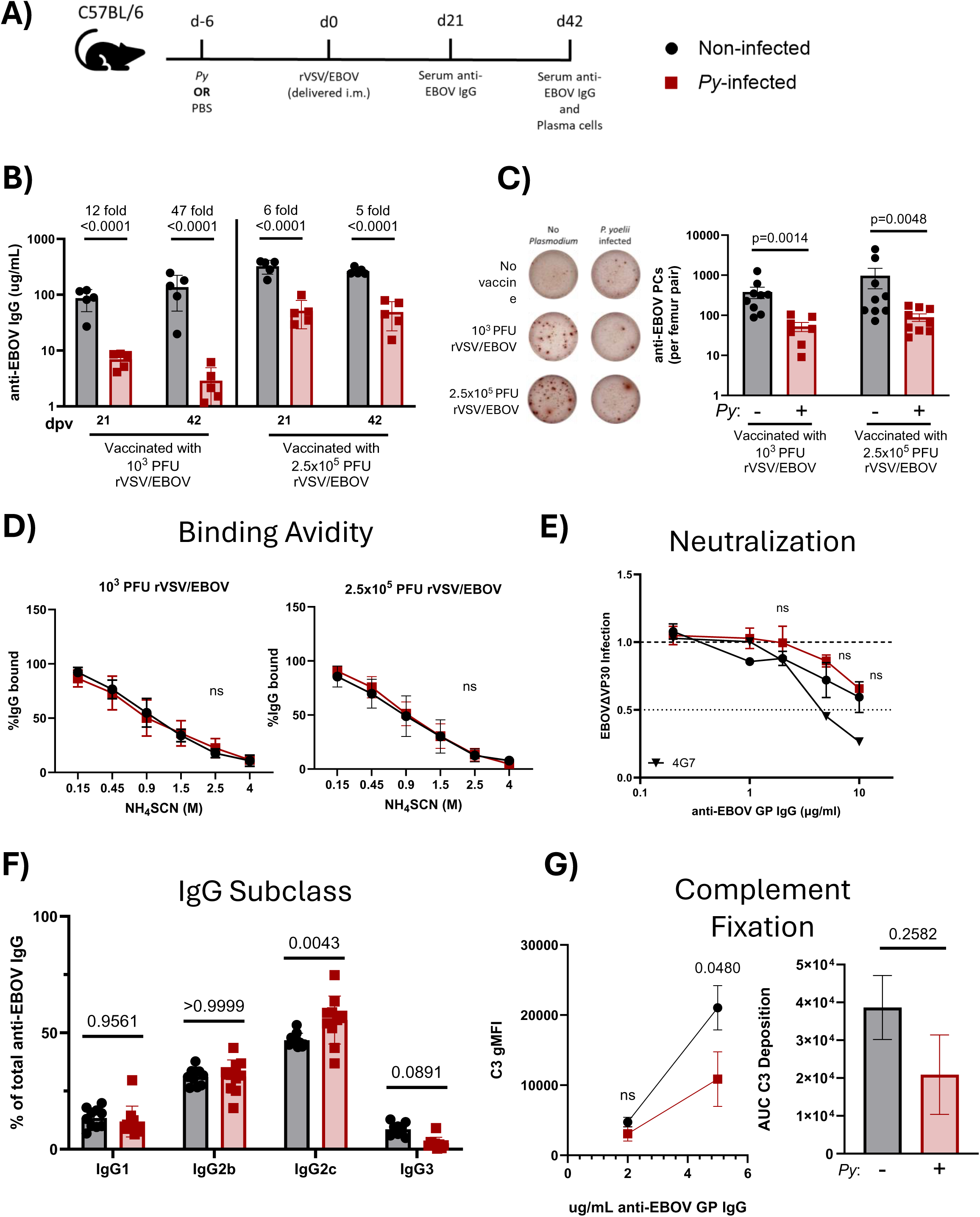
rVSV/EBOV-elicited antibodies are decreased in *Plasmodium-*infected vaccine recipients. (A) Female C57BL/6 mice were *Py*-infected (red squares) or left non-infected (black circles) and 6 days later vaccinated i.m. with 10^3^ PFU or 2.5×10^5^ PFU rVSV/EBOV. (B) Indirect ELISA probing for anti-EBOV GP IgG at 21- and 42-dpv (n=5/group). (C) ELISpot probing for anti-EBOV GP IgG-secreting cells in bone marrow at 42-dpv (n=10/group, pooled from 2 independent experiments). Representative ELISpot wells (right) and summarized cell numbers (left). (D) Indirect ELISA probing for anti-EBOV GP IgG from 42-dpv in the presence of increasing concentrations of ammonium thiocyanate (NH_4_SCN) to evaluate binding avidity. Data are represented as the percentage of binding relative to no NH_4_SCN treatment. (n=5/group, representative from 2 independent experiments). (E) Neutralization efficiency of anti-EBOV IgG derived at 42-dpv from vaccinated mice (n=3/group, representative from 2 independent experiments). Heavy dotted line represents infection of Vero VP30 cells with virus incubated with the equivalent dilution of naïve serum. Neutralization is defined as the amount of antibody that neutralizes 50% of infection, represented by the light dotted line. (F) Indirect ELISA probing for anti-EBOV IgG1, IgG2b, IgG2c, and IgG3 at 42-dpv in 2.5×10^5^ PFU rVSV/EBOV recipients. Data is represented as the percentage of IgG subclass relative to total anti-EBOV GP IgG (n=10/group, data are pooled from 2 independent experiments). (G) Bead based complement fixation of anti-EBOV IgG derived at 42-dpv from 2.5×10^5^ PFU rVSV/EBOV recipients (n=3/group, representative of 2 independent experiments). Relative gMFI of C3 binding to 2 or 5 µg/ml anti-EBOV IgG (left) and summarized AUC (right). Data in each panel were analyzed using a two-way ANOVA with Tukey’s multiple comparisons test. Data in B, D, E, and F are represented as mean ± SD. Data in C and G are represented as mean ± SEM. p<0.05 considered significant. Also see Figure S2.

In parallel studies, 2.5×10^5^ PFU rVSV/EBOV was administered intraperitoneally (i.p.) to non-infected or *Plasmodium*-infected mice at 6 days, 35 days, or 70 days following *Plasmodium* infection to examine the durability of the effect over time. We also assessed the impact of both *Py* and *P. chabaudi chabaudi* (*Pcc*) infection on rVSV/EBOV-elicited antibody responses. Infection with *Pcc* results in a less pro-inflammatory response than with *Py* but establishes chronic, recrudescing infections in mice^39^. When administered 6 days prior to vaccination, infection with *Py* or *Pcc* dramatically reduced antibody responses measured at 42-dpv **(Fig. S2C)**. The deficit caused by prior *Pcc* infection (∼11 fold) was more modest than that caused by prior *Py* infection (∼54 fold reduction). When rVSV/EBOV was administered 35 or 70 days following *Py* infection, no deficit in vaccine-elicited antibody responses was observed, indicating that the *Py-*associated mechanism(s) that suppress humoral immunity to rVSV/EBOV waned over time. While *Pcc* infection was associated with a significant increase in vaccine-elicited responses when administered 35 days prior to vaccination, a significant decrease was observed when administered 70 days prior to vaccination. This deficit imposed at 70 days following *Pcc* infection may be related to recrudescence reported with this parasite^39^.

### Functionality of EBOV GP-specific antibodies and memory-derived antibody responses are preserved in *Plasmodium-*infected rVSV/EBOV recipients

Anti-EBOV GP-specific antibodies detected at 21- and 42-dpv are associated with a primary antibody response to the vaccine, which are derived from class-switched, long-lived plasma cells (LLPC) that reside in the bone marrow. Evidence of class switched antibodies was supported by the abundance of anti-EBOV IgG at 21-dpv and undetectable levels of anti-EBOV GP-specific IgM **(Fig. S2D).** At 42 dpv, the abundance of anti-EBOV GP IgG-secreting LLPCs in the bone marrow was lower in *Py-*infected vaccine recipients compared to non-infected vaccine recipients regardless of vaccine dose **(Fig. 2C)**, consistent with a *Py-*associated deficit in the production or maintenance of vaccine-elicited LLPCs and a resulting decrease in antibody-mediated protection.

We further examined whether a *Plasmodium* infection impacts the function of EBOV GP-specific antibodies, using a series of analyses. First, we measured binding affinity of vaccine-elicited IgG to EBOV GP using an indirect ELISA with increasing concentrations of ammonium thiocyanate (NH_4_SCN). Serum antibodies derived from rVSV/EBOV-recipients at 42-dpv were bound to EBOV GP and dissociated in a dose-dependent manner with increasing concentrations of NH_4_SCN **(Fig. 2D)**, allowing evaluation of antibody affinity as previously described^40^. Binding affinity of anti-EBOV IgG derived from *Py-*infected vaccine recipients was indistinguishable from antibodies derived from non-infected vaccine recipients. Binding affinity of vaccine-elicited antibodies derived from *Py-* or *Pcc-*infected mice that had vaccine delivered i.p. rather than i.m. were similarly indistinguishable from non-infected vaccine recipients **(Fig. S2E)**, suggesting no *Py-*associated impact on binding affinity of vaccine-elicited antibodies.

Antibody neutralizing activity was next evaluated. While neutralization is not required for antibody-mediated protection against EBOV^5,7,41,42^, neutralizing antibodies are generated during rVSV/EBOV vaccination or EBOV infection and can be highly protective^5^. Because of the near undetectable levels of anti-EBOV GP IgG detected in *Py-*infected 10^3^ PFU dose recipients **(Fig 2B)**, sera derived from 2.5×10^5^ PFU recipients were utilized in the following assays. Sera derived from 42-dpv in non-infected or *Py-*infected recipients were adjusted to equivalent concentrations of anti-EBOV GP IgG and incubated with a low containment EBOV, EBOVΔVP30^43,44^ and the mixture was added to VP30-expressing Vero cells to evaluate neutralization of infectious virus. The neutralizing antibody 4G7^45^ demonstrated dose-dependent neutralization of infection and served as a positive control for these studies **(Fig. 2E)**. Modest neutralization was observed with anti-EBOV GP antibodies derived from both non-infected and *Py-*infected vaccine recipients, with no differences in neutralizing efficacy.

Functions mediated by the immunoglobulin constant region (Fc) contribute to antibody-mediated protection against filovirus infection in the context of both vaccination and therapeutic monoclonal antibodies (mAb)^7,42,46^. To evaluate whether *Py* infection results in production of different anti-EBOV IgG Fc subclasses, we measured relative amounts of IgG1, IgG2b, IgG2c, and IgG3 specific for EBOV GP at 21-dpv in both non-infected and *Py-*infected rVSV/EBOV recipients **(Fig. 2F)**. IgG2c antibodies were found to be most abundant, followed by IgG2b, IgG1, and IgG3. Anti-EBOV GP IgG derived from *Py-*infected vaccine recipients was modestly but significantly enriched in the IgG2c subclass compared to non-infected vaccine recipients.

This is not surprising, as B cell class-switching to IgG2c is driven by IFN-γ^47,48^ which is elevated during an acute *Plasmodium* infection^24–26,38,49^. Other IgG Fc subclasses were similarly represented in both non-infected and *Py-*infected vaccine recipients; however, a decreasing trend was noted for anti-EBOV GP IgG3 in *Py-*infected vaccine recipients.

The IgG3 isotype is enhanced in complement binding activity^50,51^. Given that levels of anti-EBOV GP IgG3 trended lower in *Py-*infected vaccine recipients, and activation of serum complement has been shown to play a role in antibody-mediated protection against EBOV^7,52,53^, we tested if anti-EBOV GP IgG had a different ability to bind and activate complement when derived from *Py-*infected vaccine recipients. Equivalent amounts of anti-EBOV GP IgG were assessed for their ability to activate complement via a bead-based *in vitro* assay. When EBOV GP bound beads were incubated with 5 µg/ml, but not 2 µg/ml, of anti-EBOV GP antibodies derived from *Py-*infected vaccine recipients, less C3 was bound than antibodies derived from non-infected vaccine recipients **(Fig. 2G)**. This modest difference at the higher antibody concentration suggests a potential *Py-*associated defect in the complement binding activity of anti-EBOV IgG, which may contribute to reduced vaccine-elicited protection against ma-EBOV challenge.

Although the primary LLPC-derived antibody response is most critical for the rapid protection mediated by this vaccine in an EBOV outbreak context, we sought to examine whether a secondary effector response derived from memory B cells (MBCs) was impacted similarly. To test this, non-infected or *Py-*infected 10^3^ PFU rVSV/EBOV i.m. recipients were boosted with a high dose of rVSV/EBOV to elicit a secondary effector response at 45-dpv. Anti-EBOV IgG were evaluated just prior to and 5 days following the boost (50-dpv with initial priming dose), before *de novo* antibody responses could be generated; increases in anti-EBOV antibodies during this 5-day interval can only derive from pre-existing MBCs that are expanded in response to EBOV GP exposure. Non-infected vaccine recipients had an approximate 6-fold increase in anti-EBOV GP IgG responses, while *Py-*infected vaccine recipients generated an unexpected ∼400-fold increase in anti-EBOV GP IgG to levels that were not significantly different than uninfected vaccine recipients **(Fig. S2F)**. In parallel studies, *Py-, Pcc-*infected, or non-infected mice were vaccinated with 2.5×10^5^ PFU delivered i.p. followed by a boost with the same dose at 42-dpv.

MBC-derived serum antibody responses 5 days later (47-dpv with initial priming dose) were statistically equivalent between groups initially primed during *Py* or *Pcc* infection and those primed without *Py* infection **(Fig. S2G)**. Together, these data suggest that the *Plasmodium-* imposed mechanism(s) that cause dramatic decreases in the primary LLPC-mediated antibody responses do not impact MBC-derived responses, which remain poised to respond to secondary antigen exposure.

Given the dramatic reduction in LLPC-derived anti-EBOV GP-specific antibodies caused by *Plasmodium* infection and only modest changes in antibody functionality and memory-derived responses, these data support that an acute *Plasmodium* infection is disruptive to the generation or maintenance of vaccine-induced LLPCs, but not MBCs.

### *Py* infection decreases lymph node germinal center responses to rVSV/EBOV

Given the importance of antibody responses to confer protection in EBOV outbreaks and the ability of rVSV/EBOV to rapidly elicit those responses^15,16,19,54^, we sought to understand the *Py-* driven mechanisms interfering with primary LLPC-derived antibody responses. High-affinity antibody-secreting LLPCs develop in GCs which form predominantly in secondary lymphoid tissues^55^. Following immunization, antigen exposure results in the differentiation of naïve B cells into GC B cells (GCBCs). These cells undergo somatic hypermutation and affinity maturation, eventually differentiating into LLPCs that migrate to and reside in the bone marrow, secreting antibody for years following infection or vaccination^56,57^. As anti-EBOV GP IgG-secreting LLPCs are numerically reduced in *Py-*infected vaccine recipients **(Fig. 2C)**, we examined if a concurrent infection with *Py* impacts GCBC responses. Vaccine delivery through the i.m. route permitted assessment of vaccine-specific GC responses in the draining popliteal lymph node (pLN) **(Fig. 3A, complete gating strategies in Fig. S4A-B)**. This approach provided two critical advantages: anatomic separation from blood-stage parasite-specific immune responses in the spleen, and a closer approximation of how infection impacts vaccine responses that are localized to LNs in human recipients. Maximal numbers of rVSV/EBOV-elicited GCBCs (CD19^+^B220^+^CD95^+^Bcl6^+^) were observed at 10-14 dpv in the pLN of non-infected vaccine recipients **(Fig. S3A)**. This correlated with maximal numbers of GC-localized CD4+ T follicular helper cells (GC Tfh) (CD90.2^+^CD4^+^CD44^+^PD1^hi^CXCR5^hi^) **(Fig. S3B)** that support GCBC development and differentiation^55,58^.

**Figure 3:**
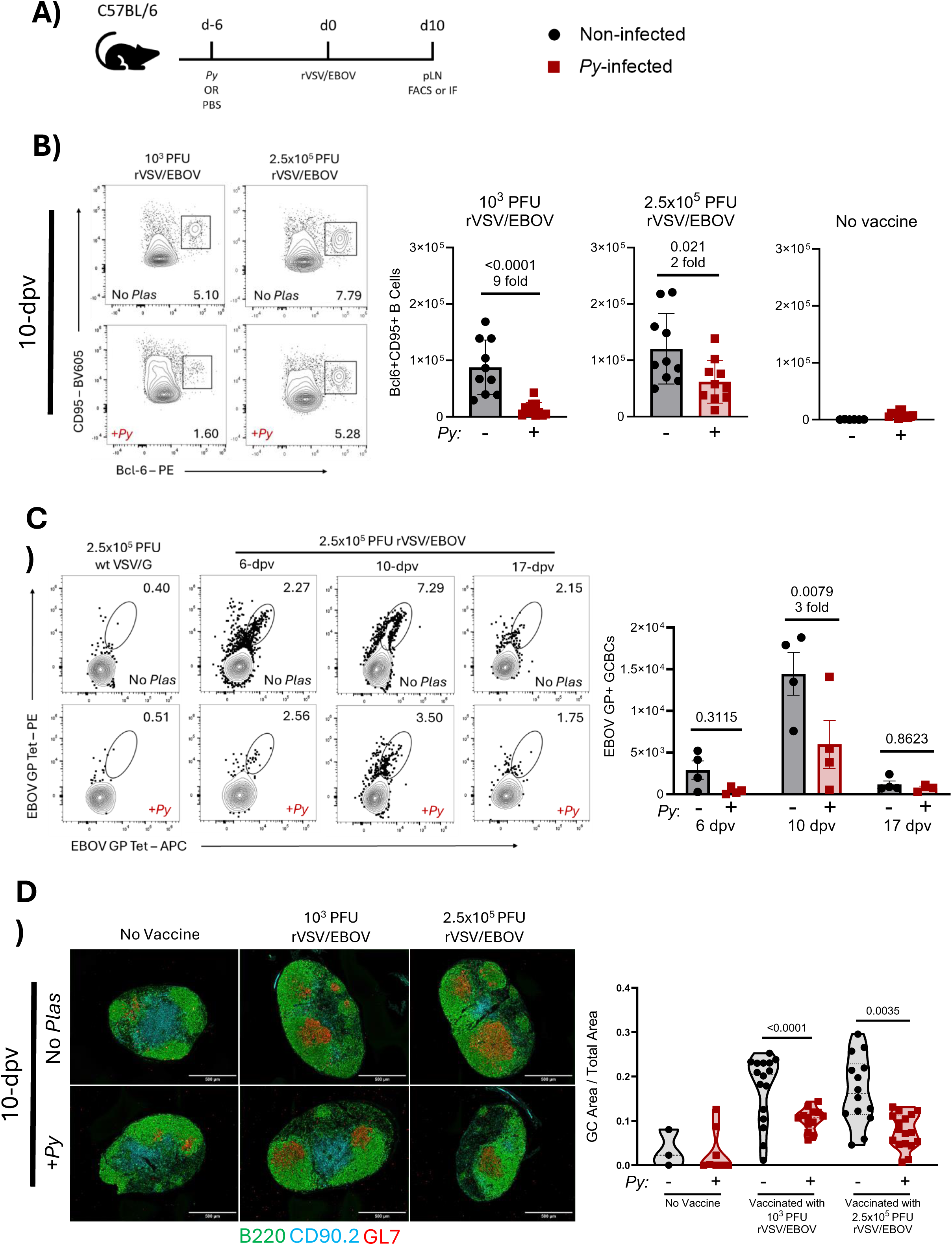
*Py* infection decreases lymph node germinal center responses to rVSV/EBOV. (A) Female C57BL/6 mice were *Py*-infected (red squares) or left non-infected (black circles) and 6 days later vaccinated i.m. with 10^3^ PFU or 2.5×10^5^ PFU rVSV/EBOV. (B) Flow cytometry evaluating GCBCs in the draining pLN at 10-dpv (n=10/group pooled from 2-3 independent experiments). Representative flow plots are gated on CD19^+^B220^+^ single cells (left) and summarized cell numbers are shown per pLN(right). (C) Flow cytometry evaluating EBOV GP-reactive GCBCs in the draining pLN of 2.5×10^5^ PFU rVSV/EBOV recipients at 6-, 10-, and 17-days post vaccination (n=4/group, one independent experiment). Representative flow plots are gated on CD19^+^B220^+^CD95^+^GL7^+^ GCBCs (left) and summarized numbers of double-positive GCBCs are shown per lymph node (right). (D) Representative images of frozen pLN sections harvested at 10-dpv (left), Original magnification, x20. Green: B220^+^ B cells; blue: CD90.2^+^ T cells; red: GL7^+^ GCBCs. Quantification of GC area relative to pLN area is shown (right) (n=2-3/group, 3-6 sections per LN, data are pooled from 2 independent experiments). Data in B were analyzed using a Student’s t-test and are represented as mean ± SD. Data in C and D were analyzed using a two-way ANOVA with Tukey’s multiple comparisons test. Data in C are represented as mean ± SEM. Violin plots in D represent the mean with bold dotted line and light dotted lines representing SD. p<0.05 considered significant. Also see Figure S3, and S4.

We examined the impact of *Py* infection on numbers of GCBCs elicited by vaccination at 10-dpv in the pLN. Non-infected recipients of 10^3^ PFU rVSV/EBOV generated ∼10^5^ GCBCs in the pLN at 10-dpv, while *Py-*infected recipients had a dramatic 9-fold decrease in numbers of GCBCs **(Fig. 3B)**, highlighting a profound impact of *Plasmodium* infection on pLN GC responses to rVSV/EBOV. A corresponding 4-fold decrease in numbers of GC Tfhs was also observed in *Py-* infected recipients at this dose **(Fig. S3C)**. GCBC numbers generated by 2.5×10^5^ PFU rVSV/EBOV were less impacted by *Py* infection, with a modest 2-fold reduction in GCBCs and no significant differences in numbers of GC Tfh **(Fig. 3B** and **S3C)**, which is consistent with the increased protection conferred by this higher vaccine dose in the presence of *Py*. Notably, *Py* infection in non-vaccinated mice elicited no appreciable GCBC responses in the pLN **(Fig. 3B)** and only modest increases in GC Tfh cells **(Fig. S3C)**, demonstrating that these responses are predominantly vaccine specific.

To resolve EBOV GP-specific responses, soluble trimeric EBOV GP lacking a transmembrane domain (EBOV GPΔTM) was conjugated to fluorescent probes to generate EBOV GP tetramers **(Fig S4C-J)**. In non-infected or *Py-*infected vaccine recipients, these tetramers identified EBOV GP-reactive GCBCs (CD19^+^B220^+^CD95^+^GL7^+^) in the pLN of rVSV/EBOV-recipients **(Fig. 3C** and **Fig S4C)**. This population was not detected in mice injected with the equivalent dose of rVSV containing its native GP (VSV/G) or mice infected with influenza A/Puerto Rico/1934 (PR8), validating its specificity **(Fig. 3C** and **S4H-J)**. A time course for production of EBOV GP-reactive GCBCs was performed on days 6, 10, and 17 following vaccinations with 2.5×10^5^ PFU **(Fig. 3C)**. While not significant, numbers of EBOV GP-reactive GCBCs trended lower in the pLNs of *Py-*infected mice at 6-dpv, suggesting that *Py* infection may impose a defect on early events in rVSV/EBOV-elicited GCBC responses. By 10-dpv, numbers of EBOV GP-reactive GCBCs were significantly decreased in *Py-*infected vaccine recipients. Few GCBCs were detected at 17-dpv in either non-infected or *Py-*infected vaccine recipients, indicating resolution of the vaccine-induced response.

Disorganized GC architecture is observed in the spleen of both humans and experimentally infected animals during *Plasmodium* infection, contributing to impaired parasite-specific antibody responses^59–61^. However, the impact of *Plasmodium* on GC architecture in LNs is not extensively studied. To evaluate GC architecture in our model, rVSV/EBOV-induced GCs were visualized via immunofluorescence microscopy (IF) at 10-dpv. As expected, substantial GC responses were observed in draining pLNs from non-infected vaccine recipients **(Fig 3D)**.

Surprisingly, pLN GCs were intact in all vaccinated mice, regardless of *Py-*infection. Quantification of GCBC-expressed GL7^+^ foci demonstrated that GC area (total and relative to LN) in draining pLNs is significantly reduced in *Py-*infected vaccine recipients compared to non-infected recipients **(Fig. 3C** and **S3D)**, while the size of the pLN and number of GCs were unchanged. **(Fig. S3D)**. Together, these findings indicate a *Plasmodium*-imposed decrease in the magnitude of rVSV/EBOV-elicited GC responses in the pLN without apparent alterations in GC architecture.

### *Py* infection induces the expression of IFN-γ-associated genes in multiple cell types in the popliteal lymph node

An acute, blood-stage *Plasmodium* infection is known to elicit systemic inflammation, including a substantial elevation of IFN-γ^25,26,62^. This impacts the composition and function of immune cells during parasite-specific humoral immune responses that take place in the spleen^63–65^. We postulated that IFN-γ may play a role in the *Py-*imposed defect in rVSV/EBOV-elicited responses in the pLN; however, how acute *Plasmodium* infection and associated elevation of IFN-γ impacts LNs is poorly understood. We sought to define the cellular profiles and transcriptional landscape of the pLN 6 days following *Plasmodium* infection to investigate changes that may contribute to poor rVSV/EBOV-elicited B cell responses.

Single cell RNA sequencing was performed on cells taken from pLNs of mice at day 0 (naïve) or day 6 following *Py* infection. To understand the impact of IFN-γ alone, we additionally performed this analysis in mice administered 3.3 µg of recombinant IFN-γ for 24 hours **(Fig. 4A).** Visualization and clustering of these samples on uniform manifold approximation and projection (UMAP) plots showed modest alterations in clustering based on treatment **(Fig. 4B)**. However, analysis of pLN cells derived from *Py-*infected or IFN-γ-treated mice showed a shared increase in expression of multiple genes, with notable increases in genes associated with IFN-γ signaling, including *Stat1, Sp100,* and *Rnf213* **(Fig. 4C)**. Further analysis of these cells identified seven distinct clusters when visualized on a UMAP plot, including large populations of B and T lymphocytes **(Fig. 4D-F)**. Small clusters corresponding to macrophages, natural killer (NK) cells, migratory dendritic cells (DCs), plasmacytoid DCs were also observed based on known markers. Proportions of these cell populations remained broadly consistent regardless of treatment, aside from a reduction of macrophages notable during both *Py* infection and IFN-γ treatment **(Fig. 4F)**. Because of low cell numbers, these smaller populations of cells were excluded from downstream analysis. We proceeded to more closely examine heterogeneity in B and T lymphocytes. We performed an isolated clustering analysis for these two subsets **(Fig. S5A, S5B).** Both B and T lymphocytes showed clusters that were divergent in *Py-*infected or IFN-γ-treated mice relative to naïve cells **(Fig. S4A, S5B)**. When applying a GSEA pathway analysis on differentially expressed genes (DEGs) in B or T lymphocytes from *Py-*infected mice compared to naïve, there was enhanced expression of genes associated with type II IFN signaling, indicating responsiveness to IFN-γ **(Fig. 4G** and **S5C, S5D)**. DEGs that had a >3 log_2_fold increased expression over cells from naïve mice were evaluated and compared between *Py* and IFN-γ recipients **(Fig. 4H)**. In B lymphocytes, there were 9 DEGs shared by both treatments, including *Gbp10, Oasl2,* and *Ifit3b*, among others **(Fig. 4H, S5E).** In T lymphocytes, there were 10 DEGs shared between treatments that again included *Gbp10, Ifit3b,* as well as *Fbxo39*, among several others **(Fig. 4H, S5F)**. In addition to shared DEGs, there were DEGs that were uniquely expressed following *Py* infection or IFN-γ treatment, indicating that some changes elicited by *Py* infection cannot be fully explained by exposure to type II IFN **(Fig. 4H)**. We used RT-qPCR to validate that at day 6 of *Py* infection, pLNs had an increased expression of *Ifng*, as well as *Irf1* and *Gbp5* **(Fig. 4I-K)**, two genes that are highly associated with IFN-γ signaling^66^. Together, these data provide evidence that an acute *Py* infection imposes a pro-inflammatory expression profile within the pLN that is enhanced in IFN-γ-responsive genes.

**Figure 4:**
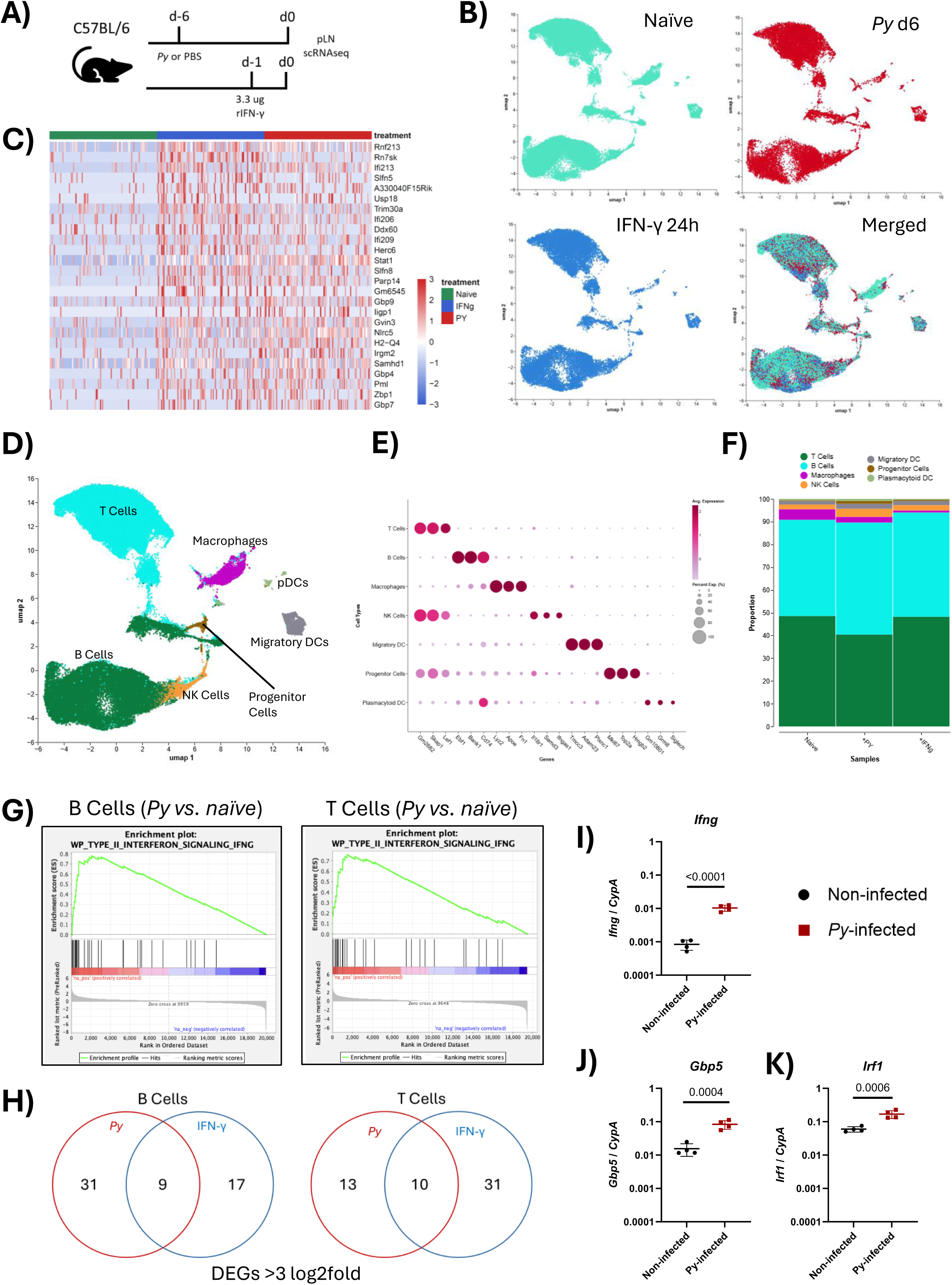
Experimental malaria induces the expression of a range of pro-inflammatory genes in multiple cell types in the popliteal lymph nodes. (A) Female C57BL/6 mice were either left non-infected, infected with Py, or treated i.p. with 3.3 μg murine recombinant IFN-γ. pLNs were harvested 6 days following Py infection or 24 hours following IFN-γ treatment and processed for single cell RNA sequencing. (B) Uniform manifold approximation and projection (UMAP) visualization of assayed samples by single cell transcriptome profiles. Clustering of individual samples alone and together is shown. (C) Heat map plot showing multiple differentially regulated genes (DEGs) in each assayed sample. (D) UMAP visualization of the cell-type composition of assayed samples by single cell transcriptomic profiles. Distinct cell types are depicted with different colors. (E) Dot plot of marker genes for distinct cell types. Color scale indicates the average normalized expression of marker genes in each cell type, and dot size is proportional to the percentage of cells within each cell cluster expressing the marker genes. Cell cluster IDs correspond to those in C. (F) Proportion of each defined cell type across groups. Color scale and size scale are the same as those in C. (G) 3 > log2fold increased DEGs over naïve compared between Py-infection and IFN-γ treatment. (H) GSEA analysis showing enrichment of genes associated with type II IFN signaling in either B (left) or T (right) cells from 6-day Py-infected mice. (I-K) RT-qPCR amplification of Ifng (I), Irf1 (J) and Gbp5 (K) from pLNs of non-infected (black circles) or 6-day Py-infected (red squares) mice. (n=4/group, data representative of 2 independent experiments). Data in I-K were analyzed using Student’s t-tests and are represented as mean ± SD. p<0.05 considered significant. See also Figure S4.

### *Py* infection causes an IFN-γ-dependent reduction in rVSV/EBOV-elicited antibody responses due in part to reduced vaccine replication

Given that the pLN in *Py-*infected mice showed transcriptional signatures consistent with IFN-γ signaling, we tested whether IFN-γ mediates the *Py-*imposed decrease in antibody responses to rVSV/EBOV. We utilized our sequential infection and vaccination model in C57BL/6 mice that lacked expression of IFN-γ (IFN-γ^-/-^). Consistent with previous reports^26,49,62^, IFN-γ^-/-^ mice infected with *Py* had a higher parasite load and delayed clearance compared to wild-type (IFN-γ^+/+^) animals **(Fig. S6A)**. Mice with or without *Py* infection were vaccinated with 10^3^ PFU rVSV/EBOV and downstream antibody responses were assessed. IFN-γ^-/-^ vaccine recipients infected with *Py* had anti-EBOV GP IgG at 21-dpv and 45-dpv that was statistically indistinguishable from non-infected recipients, in contrast with dramatic decreases caused by *Py* infection in IFN-γ^+/+^ recipients **(Fig. 5A).** This underlines a central role for parasite-elicited IFN-γ in the decreased antibody responses to rVSV/EBOV. Administration of 1 or 3.3 µg of recombinant IFN-γ i.m. 24 hours prior to vaccination recapitulated the lower anti-EBOV GP IgG levels caused by *Py* infection **(Fig 5B)**. Together, these data implicate IFN-γ as a predominant mediator of the *Py-*associated decrease in rVSV/EBOV-elicited antibody responses.

**Figure 5:**
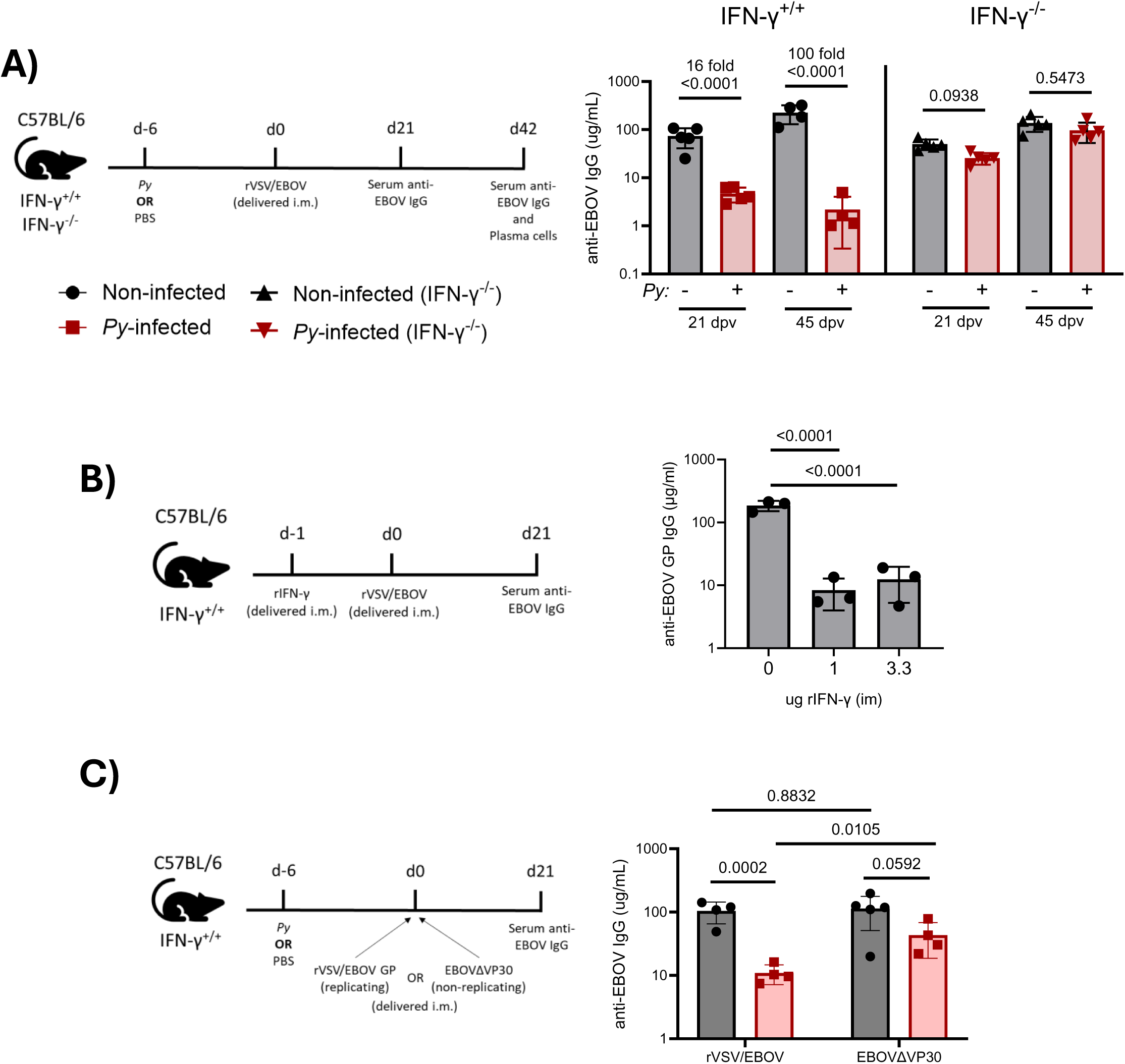
Experimental malaria causes an IFN-γ-dependent reduction rVSV/EBOV-elicited antibody responses that is partially dependent on vaccine replication. (A) Female C57BL/6 IFN-γ^+/+^ or IFN-γ^-/-^ mice were *Py*-infected (red squares or red triangles) or left non-infected (black circles or black triangles) and 6 days later vaccinated i.m. with 10^3^ PFU. Anti-EBOV GP IgG was measured at 21- or 45-dpv (n=5/group, data representative of 2 independent experiments). (B) Female C57BL/6 IFN-γ^+/+^ were treated with an i.m. injection of 0 (PBS only), 1 or 3.3 µg of murine rIFN-γ 24 hours prior to i.m. vaccination with 10^3^ PFU rVSV/EBOV in the ipsilateral leg and anti-EBOV GP IgG was measured 21 dpv (n=3/group, representative of 2 independent experiments). (C) C57BL/6 IFN-γ^+/+^ mice were Py-infected (red squares) or left non-infected (black circles) and 6 days later vaccinated i.m. with either 10^3^ PFU rVSV/EBOV i.m. OR a high dose of EBOVΔVP30 sufficient to elicit comparable IgG responses at 21-dpv, and anti-EBOV IgG was assessed at 21-dpv (n=5/group, data representative of 2 independent experiments). Data in B and D were analyzed using a two-way ANOVA with Tukey’s multiple comparisons test. Data in C was analyzed using a one-way ANOVA with Tukey’s multiple comparisons test. All data are represented as mean ± SD. p<0.05 considered significant. See also Figure S5.

Replication of rVSV/EBOV is inhibited *Py-*elicited IFN-γ signaling in peritoneal macrophages^25,66^. Given the dependency of the *Py-*imposed antibody response defect on IFN-γ, we wondered if IFN-γ-mediated inhibition of rVSV/EBOV replication contributes to *Py-*imposed deficit in antibody responses. We reasoned that if this were true, *Py* infection would have a lesser impact on immune responses elicited by the non-replicating vaccine candidate, EBOVΔVP30^43,44^. A dose of EBOVΔVP30 or UV-inactivated rVSV/EBOV with EBOV GP content equivalent to 10^3^ PFU rVSV/EBOV were both insufficient to elicit a substantial GC response **(Fig. S6B,C)**, underlining the importance of rVSV/EBOV replication to its single dose efficacy. To compare the impact of *Py* infection on immune responses to a replicating (rVSV/EBOV) versus non-replicating (EBOVΔVP30) vaccine, IFN-γ^+/+^ mice with or without *Py* infection were vaccinated with 10^3^ PFU of rVSV/EBOV or a high dose of EBOVΔVP30 that yielded equivalent levels of serum anti-EBOV IgG at 21-dpv in non-infected mice **(Fig. 5C)**. *Py-*infected recipients of EBOVΔVP30 generated significantly more anti-EBOV IgG than *Py-*infected rVSV/EBOV recipients, suggesting that *Py* infection inhibits the replication of rVSV/EBOV to mediate decreased antibody responses. Numbers of GCBCs elicited by high doses of EBOVΔVP30 or UV-inactivated rVSV/EBOV were similarly less impacted by *Py-*infection **(Fig. S6D,E)**, further highlighting that inhibition of vaccine replication is imposed by *Py* infection to result in decreased antibody responses. Notably, infection with *Py* still caused a lower trend in anti-EBOV IgG at 21-dpv in EBOVΔVP30 recipients **(Fig. 5C)**, suggesting additional *Py-*mediated mechanisms that restrict antibody responses and are independent of vaccine replication.

Together, these data indicate that IFN-γ is required for the *Py-*associated decrease in rVSV/EBOV antibody responses, which is predominantly dependent on replication-competence of the vaccine.

### *Py* infection imposes an IFN-γ-dependent decrease in rVSV/EBOV replication in the draining LN

Given that the *Py-*imposed deficit is dependent on vaccine replication, we more closely investigated the consequences of *Py* infection on rVSV/EBOV vaccine load at early time points in the draining LN. pLNs were harvested at 0.5-, 1-, 6-, 12-, or 24-hours post-vaccination (hpv) of non-infected or *Py-*infected rVSV/EBOV recipients, and vaccine load was assessed via real-time quantitative PCR (RT-qPCR) amplification of VSV matrix RNA (**Fig 6A**). These studies were done with both IFN-γ^+/+^ and IFN-γ^-/-^ mice.

**Figure 6:**
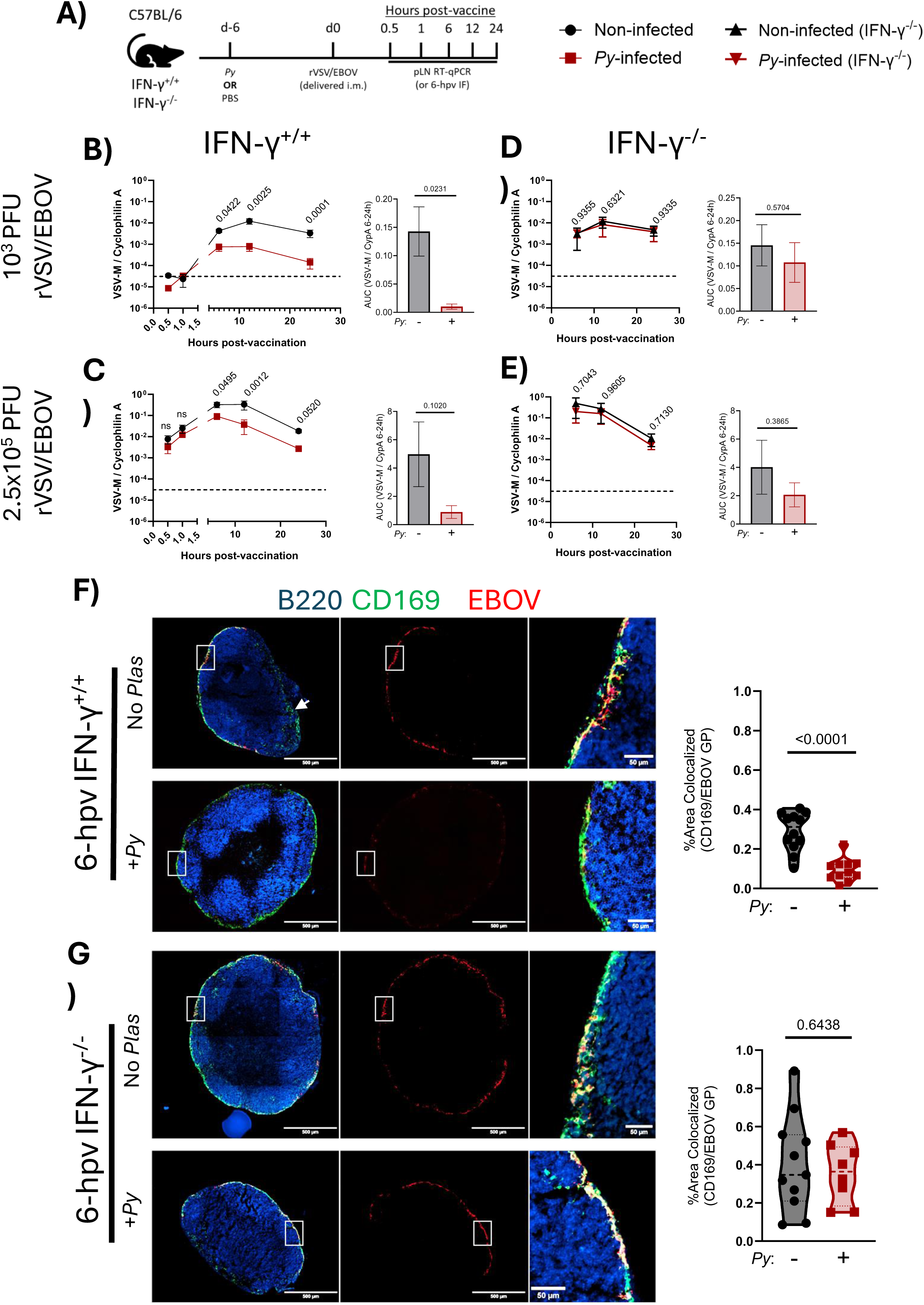
Experimental malaria imposes an IFN-γ-dependent decrease in rVSV/EBOV replication in the draining LN. (A) Female C57BL/6 IFN-γ^+/+^ or IFN-γ^-/-^ mice were either *Py*-infected (red squares or red triangles) or left non-infected (black circles or black triangles) and 6 days later vaccinated i.m. with either 10^3^ PFU or 2.5×10^5^ PFU rVSV/EBOV. Draining pLNs were harvested at 0.5-, 1-, 6-, 12-, and 24-hpv for RT-qPCR. (B, C) Amplification of VSV-M from draining pLN of 10^3^ PFU IFN-γ^+/+^ recipients (B) or 2.5×10^5^ PFU IFN-γ^+/+^ recipients (C). Summarized AUC is shown on the right (n=4/group per data point, representative from 2 independent experiments). (D, E) Amplification of VSV-M from draining pLN of 10^3^ PFU IFN-γ^-/-^ recipients (D) or 2.5×10^5^ PFU IFN-γ^-/-^ recipients (E). Summarized AUC is shown on the right (n=4/group per data point, representative from 2 independent experiments). (F, G) Representative images (left) and quantification (right) of frozen pLN sections harvested at 6 hours post vaccination with 2.5×10^5^ PFU from IFN-γ^+/+^ (F) or IFN-γ^-/-^ (G) mice. White arrow indicates hilum of the lymph node. Original magnification, x20. Green, CD169^+^ macrophages; blue, B220^+^ B cells; red, EBOV GP. Quantification of images is represented as a percentage of the area of the LN positive for both CD169 and EBOV GP (n=2-3/group, 3-6 sections per LN, data are pooled from 2 independent experiments). VSV-M RNA is normalized to expression of C*yclophilin A*. Data in B-E were analyzed using two-way ANOVA with Tukey’s multiple comparisons test and are represented as mean ± SD. Summarized AUC was analyzed via a Student’s t-test and are represented as mean ± SEM. Data in F and G were analyzed using a Student’s t-test and represent the mean with bold dotted line, with light dotted lines representing SD. p<0.05 considered significant. See also Figure S6.

In IFN-γ^+/+^ mice, administration of 10^3^ PFU did not result in detectable vaccine loads in the pLN at 0.5- and 1-hpv **(Fig. 6B)**. However, vaccine load was readily detectable at 6-hpv, indicating vaccine delivery and/or replication between 1- to 6-hpv. Vaccine load increased at 12-hpv and began to wane by 24-hpv. VSV infection is known to elicit strong type I and II interferon (IFN) responses that effectively control virus replication^67–69^, a possible explanation for the decrease in vaccine load at 24-hpv. IFN-γ^+/+^ *Py-*infected recipients had significantly lower vaccine loads at 6-, 12-, and 24-hpv, supporting that vaccine replication may be impaired by acute *Py* infection. Both IFN-γ^+/+^ recipients of 2.5×10^5^ PFU rVSV/EBOV had detectable vaccine loads at 0.5- and 1- hpv that were statistically similar between non-infected and *Py-*infected mice **(Fig. 6C)**, indicating successful delivery of this higher dose of vaccine to the pLN occurs in this time frame and is not impacted by acute *Py* infection. By 6-hpv, non-infected recipients of 2.5×10^5^ PFU had vaccine loads that were ∼100 fold higher than 10^3^ PFU recipients. Vaccine load in the pLN of *Py-*infected 2.5×10^5^ PFU recipients was also decreased between 6 and 24-hpv, perhaps due to reduced vaccine replication **(Fig. 6C)**. EBOV GP in the pLN as assessed via immunoblot at 12- hpv was also decreased in *Py-*infected vaccine recipients **(Fig. S7A)**, demonstrating that *Py-* associated changes in vaccine RNA are representative of changes in the EBOV GP antigen load. In IFN-γ^-/-^ recipients of either dose, *Py-*infection had no impact on vaccine load at 6-, 12-, and 24-hpv, when the deficit was most pronounced in IFN-γ^+/+^ mice **(Fig. 6D,E)**. These data support a central role for IFN-γ in the *Py-*imposed reduction of vaccine detected in the pLN. Notably, vaccine load kinetics from 6- to 24-hpv were not impacted by an absence of IFN-γ expression in non-infected vaccine recipients.

To further evaluate the decrease in vaccine load imposed by *Py*, we assessed cell-associated GFP expression from rVSV/EBOV-GFP in the pLN at 6-hpv by IF microscopy. To ensure sufficient sensitivity to detect infected cells via IF, a 2.5×10^5^ PFU dose was utilized in the following studies. Non-infected IFN-γ^+/+^ recipients of rVSV/EBOV-GFP had detectable GFP in the draining LN at 6-hpv **(Fig. S7B)**, which was markedly reduced in the pLNs of *Py-*infected recipients. GFP expression, which is reflective of vaccine replication, was almost exclusively restricted to the subcapsular space. This is consistent with prior studies demonstrating that subcapsular sinus macrophages (SSM) residing in this structure are uniquely permissive to rVSV/EBOV infection^70^.

We reasoned that SSMs may be directly impacted by *Py-*elicited IFN-γ, resulting in decreased viral replication. SSMs are readily identified via expression of Siglec-1 (CD169) and localization to the subcapsular sinus^70–74^. We performed co-staining of EBOV GP and CD169 at 6-hpv in the draining pLN of rVSV/EBOV recipients. In non-infected IFN-γ^+/+^ and IFN-γ^-/-^ vaccine recipients, we observed substantial co-localization of EBOV GP and CD169 by 6-hpv **(Fig. 6F,G)**, in agreement with our GFP reporter vaccine studies. Notably, CD169^+^ medullary sinus macrophages (MSM) located near the hilum of the pLN did not co-stain for EBOV GP **(Fig. 6F, white arrow)**. In *Py*-infected IFN-γ^+/+^ vaccine recipients, there was a marked reduction in EBOV GP staining and in co-localization with SSMs **(Fig. 6F)**. The loss of EBOV GP was not observed in *Py-*infected IFN-γ^-/-^ mice **(Fig. 6G)**, indicating that IFN-γ mediates the *Py-*imposed decrease in rVSV/EBOV infection of SSMs. Loss of IFN-γ expression caused no apparent visual changes in SSM numbers or localization **(Fig. S7C).** Notably, *Py* infection caused a modest increase in CD169 staining in the pLN **(Fig. S7C)**, suggesting increased survival of these cells at 6-hpv in *Py-*infected vaccine recipients. SSMs are poised to undergo cell death shortly after encountering immunogenic antigen or virus^71,72,75,76^. Thus, an increase in CD169 detection may reflect a *Py-*dependent decrease in SSM cell death due to vaccine replication.

Together with our observation that a non-replicating vaccine is less susceptible than rVSV/EBOV to the *Py-*imposed antibody deficit **(Fig. 5)**, these data demonstrate that *Py* infection imposes an IFN-γ-dependent restriction of rVSV/EBOV replication in SSMs in the pLN. This decrease is associated with a reduced magnitude of primary anti-EBOV IgG responses that are critical for vaccine-mediated protection against ma-EBOV challenge.

## DISCUSSION

Given the severity of EVD, the lack of readily accessible therapeutics, and the requirement of an EBOV vaccine to provide rapid protection in emergency settings, identifying the most effective vaccine protocols are essential for the optimal management of EVD outbreaks. We show that acute, blood-stage murine *Plasmodium* infections negatively impacts the primary antibody response elicited by low dose rVSV/EBOV vaccination and results in reduced protection against a lethal dose of ma-EBOV. We defined the mechanism of this defect, showing that orchestration of vaccine-specific GCBC responses within the dLN is reduced in *Py-*infected mice. The reduction in GCBCs is largely dependent on an elevation of IFN-γ elicited by *Py* infection that restricts replication of rVSV/EBOV in SSMs, resulting in suboptimal rVSV/EBOV-driven B cell responses. Despite the reduced levels of primary rVSV/EBOV-elicited antibodies during acute *Plasmodium* infection, we observed little impact on functionality of the antibodies generated and no impact on memory-derived antibody responses upon secondary antigen exposure. Infection with *Pcc* also imposed a decrease in rVSV/EBOV-elicited antibody responses, supporting that this impact extends to other murine *Plasmodium* parasites. The *Py-*imposed decrease in vaccine-mediated protection can be overcome with higher doses of rVSV/EBOV. While the strong protection conferred by rVSV/EBOV and significant side effects known to be associated with this vaccine have led to the suggestion that the vaccine dosage be reduced^19^, our studies provide a rationale for maintaining a higher dose.

A prior study that examined the impact of asymptomatic malaria on rVSV/EBOV efficacy found only modest decreases in neutralizing vaccine-elicited antibodies up to a year following vaccination, and levels of EBOV GP-specific antibodies that were classified as protective^37^. This supports a high efficacy of the currently licensed dose even in *Plasmodium-*infected recipients and is consistent with our findings that protection mediated by high dose rVSV/EBOV is not impacted by *Py* infection. However, it should be noted that the asymptomatic malaria in participants of this study is likely reflective of exposure to repeated *Plasmodium* infections, which induces a more immunomodulatory profile^77^. Symptomatic parasitemia that is associated with a pro-inflammatory response occurs in naïve hosts, such as young children, traveling healthcare workers or tourists ^24^. It is these groups that our study models, hence, the insights provided by these studies reflect only one portion of the population requiring vaccination during EVD outbreaks. Future studies using a murine model of sequential, repeated *Plasmodium* infections can better evaluate how the unique immunological environment elicited by repeated *Plasmodium* infections impacts vaccine responses.

Other studies have examined the impact of asymptomatic malaria and clinical episodes of malaria on the two-dose regimen of Ad26.EBOV and MVA-BN-Filo that is licensed for prophylactic use in areas at risk for EVD outbreaks, finding no significant difference in antibody responses following the second dose in malaria-infected or malaria-negative recipients of any age group^78,79^. Because this is a replication-incompetent vaccine platform, the absence of any malaria-associated impact is consistent with our data using replication-incompetent EBOVΔVP30. Another probable contributor to the high efficacy of this two-dose regimen in malaria-infected individuals is that the second dose likely stimulates protective MBC-derived antibody responses, even when recipients are poorly primed during acute *Plasmodium* infection. MBC-derived antibody responses were unaffected by prior *Py* infection in our model, further supporting that a secondary dose can overcome the *Plasmodium*-associated deficit and stimulate protective levels of antibodies. Nevertheless, a notable logistical problem with a two-dose filovirus vaccine regime is the unpredictable environment during an EVD outbreak. This makes verification of a second vaccine dose problematic and may not be a feasible solution to *Plasmodium-*associated decreases in vaccine efficacy.

Due to the substantial disease burden imposed by malaria, extensive efforts to understand the mechanistic relationship between *Plasmodium* infection and heterologous vaccine effectiveness are ongoing. Our findings demonstrate that IFN-γ-dependent inhibition of rVSV/EBOV replication predominantly underlies the decreased antibody responses; however, IFN-γ is a multifunctional cytokine that plays multiple established roles in malaria immunopathology. IFN-γ elicited by murine *Plasmodium* infection impacts the differentiation of splenic T follicular helper cells in both parasite-specific responses or responses to a heterologous immunogen, resulting in reduced humoral immunity^63,64,80^*. Plasmodium*-elicited IFN-γ also directly impacts B cell function^65^. This direct impact of IFN-γ on B and T cell function likely causes additional decreases in antibody responses to rVSV/EBOV that are not dependent on vaccine replication, which is supported by a remaining modest impact of *Plasmodium* on responses to non-replicating EBOVΔVP30 or high doses of rVSV/EBOV. The multifunctional role of IFN-γ in our system is further underscored by the observed increase in frequency of anti-EBOV GP IgG2c in *Py-*infected recipients, likely due to IFN-γ-driven class-switch recombination (CSR) to the IgG2c isotype in C57BL/6 mice^47,48^.

Malaria-driven disruption of splenic GC architecture has also been implicated as an important feature leading to suboptimal antibody responses^59–61,63^. We show that this feature does not extend to the LN, as intact GCs were observed in draining pLNs of vaccine recipients regardless of *Py* infection status. Substantial architectural changes in the spleen during *Plasmodium* infection are likely driven by continuous hematogenous exposure to parasite-derived antigens and metabolites that influence the function of immune cells responding to both parasite-derived and unrelated antigens^24,49,76,81,82^. LNs may not be subject to this level of exposure, resulting in preserved GC architecture. More work must be done to more carefully determine the differential impact of malaria on LN-localized versus splenic immune responses.

Our surprising observation that MBC-derived antibody responses are preserved in mice that were primed during *Plasmodium* infection suggests several possibilities. MBCs differentiate from the GC earlier than LLPCs, with lower affinity for their target antigens^83,84^. A lower threshold for antigen affinity might suggest that the production of MBCs is less dependent on antigen dose than production of LLPCs, although this has not been shown. This would explain a preserved production of EBOV GP-specific MBCs in the context of reduced rVSV/EBOV replication. Another plausible mechanism is the extrafollicular MBC formation that occurs in response to some antigens^85^, which could serve as a source of rVSV/EBOV-elicited MBCs that are unaffected by the reduced GC responses imposed by *Plasmodium*. In our model, it is likely that the preserved MBC-derived antibody response contributes to protection (∼70%) of *Py*-infected low dose vaccine recipients against ma-EBOV infection even with low serum anti-EBOV GP antibody responses (∼10 µg/ml) that are insufficient for protection^6^. A more careful phenotypic analysis of MBCs generated in the presence or absence of a concurrent *Plasmodium* infection could provide insight into the influence of antigen dose and *Plasmodium-* elicited inflammation on GCBC output.

SSMs that reside in peripheral lymph nodes are appreciated to sequester and directly initiate B cell activation to a range of different immunogens^70,72–76,86^. We show that SSMs residing in peripheral lymph node support rVSV/EBOV replication within hours following vaccination, extending previous studies that show replication in this cell type contributes to the immunogenicity of rVSV/EBOV^70^. SSMs are uniquely permissive to infection with rVSV/EBOV and other viruses due to a reduced ability to respond to type I IFNs^70–72,87^; however, the ability of SSMs to respond to type II IFN is poorly studied. Our findings provide evidence that these cells respond to IFN-γ, decreasing rVSV/EBOV replication in the presence of an acute *Plasmodium* infection. Interestingly, the *Plasmodium-*associated elevation of IFN-γ that results in suboptimal antibody responses is also mechanistically responsible for restriction of EBOV replication in peritoneal macrophages that protects against EVD in a mouse model^25^. This suggests that, paradoxically, an acute infection with *Plasmodium spp.* may still contribute to protection against EVD even if there is a concomitant restriction of vaccine-mediated protection.

While these studies mechanistically investigate the interaction of *Plasmodium* and rVSV/EBOV, there are several associated limitations. Murine models of *Plasmodium*, while valuable tools for immunological investigation, do not perfectly recapitulate the inflammatory environment elicited by *P. falciparum* infections in immune-experienced humans. Further, rVSV/EBOV dosing in this study serves as only an approximation of relative low doses suggested for human recipients during outbreaks. Controlled investigation of the impact of *Plasmodium* infections on rVSV/EBOV efficacy in humans or NHPs are warranted to expand the clinical relevance of our findings.

Due to the remarkable efficacy of rVSV/EBOV and the proven utility of rVSV-vectors in eliciting strong immune responses, analogous rVSV-vectored vaccines expressing GPs from other filoviruses and viruses of concern are in varying stages of development^10,88–92^. Given our findings, consideration of a malaria-associated impact on rVSV-vectored vaccine efficacy will be important for optimal design of effective vaccine protocols in malaria-endemic areas. Based on these studies, we suggest that maintaining rVSV/EBOV doses that deliver high antigen loads is a promising strategy to prevent *Plasmodium-*associated deficits that could otherwise limit the efficient control of future EVD outbreaks.

## MATERIALS AND METHODS

### Ethics Statement

All work involving infectious ma-EBOV was performed following standard operating procedures (SOPs) approved by the Rocky Mountain Laboratories (RML) Institutional Biosafety Committee (IBC) in the maximum containment laboratory at the RML, Division of Intramural Research, National Institute of Allergy and Infectious Diseases (NIAID), National Institutes of Health (NIH). Mouse work was performed in strict accordance with the recommendations described in the Guide for the Care and Use of Laboratory Animals of the National Institute of Health, the Office of Animal Welfare, and the United States Department of Agriculture and was approved by the RML Institutional Animal Care and Use Committee (IACUC). All animals were housed under controlled conditions of humidity, temperature, and light (12-h light:12-h dark cycles). Food (commercial chow) and water were available *ad libitum*.

### Study Design

#### ma-EBOV studies

Assuming that 100% of mice survive in any vaccine group in the absence of malaria and 40% in the presence of malaria, groups of 10 mice were needed to achieve statistical significance with 80% power with the Fisher exact test, one-sided test alpha set to 0.025. Five mice in each necropsy group are needed to achieve statistical significance with 80% power of differences in viral loads if we assume a 1-2 log reduction in 95% of the non-infected vaccinated mice compared to *Py-*infected vaccinated mice. Animals were randomly assigned to groups by generating a randomized excel sheet of experimental groups.

#### All other studies

Samples sizes were determined based on preliminary data effect size and standard deviation metrics using a type I error rate of α=0.05 and power=80%. Preliminary power analysis suggested n>3 for each mouse experiment. In most experiments, we used a larger *n* for reliable conclusions. Sample collection was performed at pre-determined time points followed by data collection, and data were excluded when there were indications of artifact or experimental failures, such as infection failure or mouse injury. The exclusion criteria were predetermined, and no outliers were excluded. Randomization principles were used for all mouse experiments.

### Mice

C57BL/6J wild-type and B6.129S7-*Ifng^tm1Ts^*/J (IFN-γ KO) mice were purchased from The Jackson Laboratory (strain #000664, strain #002287, respectively). The University of Iowa (UI) Institutional Assurance Number is #A3021-01. All mouse procedures performed at the UI were approved by the UI Institutional Animal Care and Use Committee (IACUC) which oversees the administration of the IACUC protocols, and the study was performed in accordance with the IACUC guidelines (Protocol #1031280 and #4021280).

### Viral stocks

Ma-EBOV^93^ was propagated on Vero E6 cells, titered on these cells, and stored in liquid nitrogen. Recombinant VSV encoding the EBOV GP (rVSV/EBOV Kikwit, rVSV/EBOV-eGFP Mayinga) or the native VSV glycoprotein (rVSV/G) and EBOVΔVP30-eGFP (Mayinga) were generated as previously described ^94^. Briefly, virus was propagated by infection of Vero E6 or Vero VP30 cells at a low MOI (∼0.005) and collection of supernatants at 48 hpi for rVSV or at 120 hours for EBOVΔVP30, when robust GFP expression was observed. Supernatant was filtered through a 45-micron filter and purified by ultracentrifugation (28,000 x g, 4°C, 2 hr) through a 20% sucrose cushion. The pellet was resuspended in a small volume of sterile PBS, aliquoted and stored at -80°C until use. Viral titers were determined by plaque forming assay on Vero E6 or (tissue culture infectious dose 50%) TCID_50_ on Vero VP30 cells.

### Infections, Vaccinations and Treatments

Ma-EBOV infection was performed by diluting in 1X PBS to 10 focus forming units (FFU) per 0.2 mL immediately before challenge. Mice were inoculated intraperitoneally by injecting 0.1 mL of suspension into 2 sites of the lower abdomen as previously described ^95^. *Plasmodium yoelii* (clone 17NXL, obtained from MR4, ATCC) infections were initiated by serial transfer of 200 µl 1X PBS containing 10^6^ parasitized red blood cells (pRBCs) via intravenous injection. Vaccine (rVSV/EBOV or EBOVΔVP30) was administered at the indicated dose in 100µl 1X PBS via intramuscular injection in the anterior tibialis muscle. Similar amounts of virus were administered via intraperitoneal injection in some studies. For experiments utilizing recombinant IFN-γ (Cat: 575306, Biolegend, San Diego CA), cytokine was delivered either i.p. or i.m. at the indicated dose and time suspended in 100 µl 1X PBS.

### Titration of ma-EBOV

For determination of ma-EBOV titers in mouse blood and tissue samples, Vero E6 cells were seeded in 48-well plates the day before titration. Tissues were homogenized in 1 mL plain DMEM and tissue and blood samples were serial diluted 10-fold. Media was removed from cells and triplicates were inoculated with each dilution. After one hour, DMEM supplemented with 2% FBS, penicillin/streptomycin and L-glutamine was added and incubated at 37°C. Cells were monitored for cytopathic effect (CPE) and the TCID_50_ was calculated for each sample employing the Reed and Muench method^96^.

### Parasite Quantification

Parasitemia was measured using flow cytometry using an approach similar to previously described methods^97^. Briefly, 1 µl of blood was obtained through a tail tipping method and mixed in 30 µl of PBS. Cells were spun down and resuspended in 100 µl of a cocktail containing Hoechst-34580 (1:1000, Thermo Fisher Scientific, Waltham MA), Dihydroethidium (1:500; Thermo Fisher Scientific), CD45.2-FITC (1:200 of clone 104, Biolegend) and Ter-119-APC (1:500 of clone TER-119, Tonbo Biosciences) and incubated for 30 minutes at 4°C. The cells were washed with PBS and spun down at 200 g for 5 minutes and run on an Aurora Cytek after resuspending in 100 µl of PBS. Percentage of Ter119^+^CD45.2^-^ RBCs positive for both Hoechst-34580 and Dihydroethidium indicative of parasitized cells were calculated using FlowJo analysis software (Tree Star Inc., San Carlos CA).

### Enzyme-linked immunosorbent assay (ELISA)

Detection of anti-EBOV GP antibodies was performed by coating optical microtiter plates (Immulon®) overnight with soluble EBOV GP (Kikwit-95, Acro Biosystems, Newark DE) (50 µl/well at 1 µg/ml) or a standard curve composed of mouse Ig (Southern Biotech, Birmingham AL). Wells were washed 3x with ELISA Wash Buffer (1X PBS + 0.015% Tween 20), blocked for 2 hours at 25°C with ELISA Blocking Buffer (1X PBS + 2% BSA; 200 µl/well) and incubated overnight at 4°C with serum obtained from immunized mice. Following incubation, wells were washed as described and incubated with horseradish peroxidase (HRP)-conjugated anti-mouse IgG/M, IgG, IgG1, IgG2c, or IgG3 (1:500 in 50 µl ELISA blocking buffer, Jackson ImmunoResearch Inc., West Grove PA) for 1 hour at 25°C. Wells were washed as described and incubated with HRP substrate according to the manufacturer’s protocol (BD Biosciences, Milpitas CA). The reaction was stopped with 50 µl 2M H_2_SO_4_ and absorbance at 450 nm was measured on a Tecan Plate Reader (Life Sciences, Mannedorf Switzerland). For IgG and IgG/M assays, total concentration of anti-EBOV GP antibodies was quantified via comparison to the linear range of the standard curve. For IgG subtypes, the absorbance of each subtype was added together to obtain a total concentration, which was then used to determine the percentage of each subtype out of total anti-EBOV GP IgG. To assess binding avidity, anti-EBOV GP Ig from serum of immunized mice was adjusted to comparable concentrations, bound to EBOV GP, and incubated with NH_4_SCN for 15 minutes before addition of HRP-conjugated anti-mouse IgG^40^. Binding avidity is evaluated as the concentration of NH_4_SCN required to dissociate 50% of anti-EBOV GP measured via absorbance at 450 nm.

### Enzyme-linked immunosorbent spot (ELISpot)

White polystyrene plates (Nunc Maxisorp, Nunc-Immuno) were coated with soluble EBOV GP (Kikwit-95, Acro Biosystems, Newark DE) (50 µl/well at 2 ug/ml). Plates were washed with 1X PBS and blocked in ELISA Blocking Buffer for 2 hours at 25°C. Bone marrow cells isolated from immunized mice were counted and serially diluted in Roswell Park Memorial Institute (RPMI) 1640 Medium supplemented with antibiotics and 10% Fetal Bovine Serum (FBS) and 200 ul added to each well. Cells were incubated for 18 hours at 37°C with 5% CO_2_. Following extensive washes with ELISA Wash Buffer, HRP-conjugated anti-mouse IgG was added for 1 hour at 25°C (diluted in ELISA Blocking Buffer, Jackson ImmunoResearch Inc.). Following extensive washes, spots were developed using 3-amino-9-ethylcarbazole. Spots were quantified using an automated ELISpot counter.

### Neutralization Assay

For assessment of neutralizing efficiency of vaccine-elicited anti-EBOV GP antibodies, heat-inactivated serum from immunized mice was adjusted to equivalent concentrations and incubated with the surrogate EBOV model, EBOVΔVP30 ^43^, for 1 hour at 37°C. To control for antibody-independent serum inhibitors of virus infection, equivalent dilutions were made with serum from non-immune mice. After incubation, virus was added to Vero VP30 cells at an MOI of 1 for 24 hpi, then assessed for the % of GFP+ infected cells on a CytoFLEX flow cytometer (Beckman, Brea CA). 100% infectivity was defined as % of infected cells when virus was incubated with an equivalent dilution of non-immune serum, and neutralization is defined here as the concentration of anti-EBOV GP IgG sufficient to reduce 50% or more of virus infectivity.

### Flow Cytometry

Popliteal lymph nodes were forced through a 70-µm mesh to generate single-cell suspensions. Single cell suspensions were subjected to RBC lysis, counted, then resuspended at a cell density of 10^6^ cells per 100 µl. Cell-specific staining protocols were utilized as described below, using antibodies diluted in FACS buffer (PBS + 0.09% sodium azide + 2% fetal bovine serum). For detection of GCBCs, single cells were stained with an antibody cocktail consisting of Fc block/CD16/32 (clone 2.4G2) and the following α-mouse antibodies: CD19-APC-Cy7 (1:300 of clone 6D5, Biolegend), CD45R/B220-AF700 (1:300 of clone RA3-6B2, Biolegend), CD38-APC (1:300 of clone 90, Biolegend), Fas/CD95-BV605 (1:300 of clone SA367H8, Biolegend). Cells were washed in FACS buffer, spun at 700 x *g* for 5 minutes at 4°C and resuspended in foxp3/transcription factor fixation/permeabilization buffer (Tonbo Biosciences) for 15 minutes at room temperature. Cells were then spun at 700 x *g* for 5 minutes at 4°C and resuspended in Flow Cytometry Perm Buffer (Tonbo Biosciences) containing α-mouse Bcl-6 (1:100 of clone K112-91, BD Biosciences). For detection of GC Tfh cells, single cells were stained with an antibody cocktail consisting of Fc block/CD16/32 (clone 2.4G2) and the following α-mouse antibodies: CD90.2-AF647 (1:2000 of clone 30-H12, Biolegend), CD4-PerCP-Cy5.5 (1:300 of clone GK1.5, Biolegend), CD44-BV510 (1:100 of clone IM7, Biolegend), PD1-PE-Cy7 (1:200 of clone RMP1-30, Biolegend), and CXCR5-BV421 (1:100 of clone L138D7, Biolegend). Following staining protocols, cells were then evaluated on an Aurora Cytek Flow Cytometer (Aurora). Enumeration of cells via flow cytometry was performed by multiplying the frequency of indicated cell population out of live cells by the total number of live cells obtained from the LN.

### Antibody-Dependent Complement Deposition (ADCD)

To evaluate the ability of anti-EBOV GP antibodies to bind and activate complement, EBOV GP (Kikwit-95, Acro Biosystems) was biotinylated using LC-LC-Sulfo-NHS Biotin (Thermo Fisher Scientific). Excess biotin was removed using a Zeba desalting column (Thermo Fisher Scientific). Biotinylated GP antigen was coupled to 1 um red Neutravidin beads (Thermo Fisher Scientific) via overnight incubation at 4°C. Beads were washed 2x with 0.1% BSA. Serum from immunized mice was diluted in unsupplemented RPMI to 5 µg/ml or 2 µg/ml anti-EBOV GP as determined by ELISA, and incubated with GP-coupled beads for 2 hours at 37°C. Beads were washed 3x with ELISA Wash Buffer to remove unbound antibodies prior to the addition of reconstituted guinea pig complement (Cedarlane Labs, Burlington ON) diluted in veronal buffer supplemented with calcium and magnesium (Boston Bioproducts, Milford MA) for 20 minutes at 37°C. Beads were washed with PBS containing 15 mM EDTA and stained with a Fluorescein-5-isothiocyanate (FITC)-conjugated anti-guinea pig C3 antibody (MP Biomedicals, Santa Ana, CA). C3 deposition onto beads was measured using a CytoFLEX flow cytometer (Beckman). The geometric mean fluorescent intensity (gMFI) of FITC on fluorescent beads was measured, and data is represented as Area Under Curve (AUC) calculated from gMFI of beads incubated with varying concentrations of anti-EBOV IgG.

### Immunofluorescence Imaging

To evaluate pLN microarchitecture, pLNs from immunized mice were fixed in 4% paraformaldehype (PFA) for 24 hours, dehydrated in 30% sucrose for at least 48 hours, then embedded in optimal cutting temperature (OCT) freezing media and frozen using dry ice. Tissue blocks were cryosectioned at 10 um thickness using cryostat (Thermo Fisher). Mounted sections were washed with 1X PBS to remove residual OCT media, then blocked for 2 hours at 25°C using an IF blocking buffer (1X PBS + 5% BSA + 5% Normal Goat Serum + 0.1% Fish Skin Gelatin). For staining germinal centers, sections were stained overnight at 4°C using antibodies diluted in IF blocking buffer; anti-mouse B220-AF488 (1:100 of clone RA3-6B2, Biologend), anti-mouse/human GL7 Antigen-AF647 (1:30 of clone GL7, Biolegend), and anti-mouse CD90.2-biotin (1:1000 of clone 53-2.1, Biolegend). Sections were then washed 3x for 5 minutes with wash buffer (1X PBS + 0.015% Tween 20) and counter stained with streptavidin-AF546 for 1 hour at 25°C. For staining EBOV GP, sections were stained overnight at 4°C using antibodies diluted in IF blocking buffer; anti-mouse B220-eFluor450 (1:100 of clone RA3-6B2; eBioscience), anti-mouse CD169-AF488 (1:50 of clone 3D6.112; Biolegend), and human anti-EBOV GP (1:100 of clone KZ52, IBT Bioservices, Rockville MD). Sections were then washed 3x for 5 minutes with wash buffer (1X PBS + 0.015% Tween 20) and counter stained with anti-human AF647 (Jackson ImmunoResearch for 1 hour at 25°C. For staining GFP, sections were stained overnight at 4°C using anti-GFP-AF488 (1:50; Invitrogen).

Sections were finally washed 3x for 5 minutes with wash buffer (1X PBS + 0.015% Tween 20), overlaid with ProLong Diamond Antifade Mountant (ThermoFisher Scientific) and left to set for 24 hours. Sections were visualized using a Zeiss microscope at indicated magnification and images were processed using Zeiss Blue Software (Oberkochen, Germany). Analysis of total lymph node area and germinal center area was performed by manually tracing the lymph node border and GL7+ foci. EBOV GP co-localization was calculated by manually adjusting the threshold and calculating the percentage of pixel area in each section co-stained for EBOV GP and CD169. All image analysis was performed using ImageJ (NIH, Bethesda MD)

### Single-Cell RNA Sequencing

Popliteal lymph nodes were forced through a 70-µm mesh to generate single-cell suspensions. Cells were pooled from multiple lymph nodes from day 0 (naïve), day 6 following *Py* infection, or 24 hours following intraperitoneal injection of 3.3 µg recombinant IFN-γ. Cells were fixed and processed for sequencing using the Evercode WT single-cell whole-transcriptome kit (Parse Biosciences, Seattle WA) according to the manufacturer’s guidelines. Resulting nucleic acid sublibraries were evaluated for adequacy on an Agilent Bioanalyzer (RNA integrity number >8), pooled, and sequenced on an Illumina Novaseq 6000 sequencer using 100 base pair (bp) paired-end reads (Iowa Institute of Human Genetics, University of Iowa). Resulting FASTQ files were processed using the Parse Biosciences pipeline (v1.1.1) with default settings to align sequencing reads to the mouse genome and generate gene-cell count matrices. Downstream processing and analysis were performed using the Parse Trailmaker software and the R package Seurat (v5.0.0). After excluding low-quality cells, potential doublets, and dead cells, we obtained a total of 104,454 cells, composed of 60,028 (Day 0), 20,416 (Day 6 *Py*), and 24,010 (IFN-γ) across one biological replicate. Gene set enrichment analysis (GSEA) plots were generated using the GSEA software (MIT, MA).

### RNA isolation and RT-qPCR

To detect viral RNA in the lymph nodes and spleens of immunized mice, tissue was harvested from mice at time points indicated in figure legends. Harvested tissue was immediately homogenized in TRIzol reagent (Invitrogen) followed by RNA extraction according to the manufacturer’s specifications. 1 µg of RNA from each sample was converted to cDNA using a High-Capacity cDNA RevTrans Kit (Applied Biosystems). Quantitative PCR was performed using POWER SYBR Green Master Mix (Applied Biosystems) according to the manufacturer’s specifications. Data were collected on a QuantStudio 3 Real Time PCR instrument and Ct values determined with the QuantStudio Data Analysis software (Applied Biosystems).

Averages from duplicate wells for each gene were used to calculate abundance of transcripts relative to the housekeeping gene (c*yclophilin A*) and represented as 2^-ΔCt^. Primers were obtained (Integrated DNA Technologies, Coralville IA) and sequences are as follows; VSV-M fwd 5’-CCTGGATTCTATCAGCCACTT-3’, VSV-M rev 5’-TTGTTCGAGAGGCTGGAATTA-3’, *Ifng* fwd 5’-CGGCACAGTCATTGAAAGCC-3’, *Ifng* rev 5’-TGTCACCATCCTTTTGCCAGT-3’, *Gbp5* fwd 5’-CCCAGGAAGAGGCTGATAG-3’, *Gbp5* rev 5’-TCTACGGTGGTGGTTCATTT-3’, *Irf1* fwd 5’-TCACACAGGCCGATACAAAG-3’, *Irf1* rev 5’-GATCCTTCACTTCCTCGATGTC-3’, *CypA* fwd 5’-GCTGGACCAAACACAAACGG-3’, *CypA* rev 5’-ATGCTTGCCATCCAGCCATT-3’.

### Generation of EBOV GP Tetramers

FreeStyle™ 293-F Cells (ThermoFisher Scientific) were co-transfected with plasmids encoding EBOV GP-6XHIS and EBOV GP-AviTag at a 2:1 ratio using 293fectin Reagent (Gibco). Transfected cells were cultured in FreeStyleTM 293 Expression Medium for three days and the supernatant was collected by centrifugation at 4,000 x g for 30 minutes. Recombinant EBOV GP was purified by FPLC using a HisTrap HP Column (Cytiva) and eluted with 250 mM of imidazole. The purified EBOV GP was biotinylated using BirA biotin-protein ligase kit (Avidity). Protein purification was confirmed by SDS-PAGE **(Fig. S8A)**. Biotinylated EBOV GP was tetramerized with fluorochrome-labeled streptavidin (Agilent). Labeled tetramers were centrifuged at 20,000 x g for 10 minutes to remove aggregates and precipitates. To validate the tetramers, three types of mediastinal lymph node cells were stained with EBOV GP-PE and EBOV GP-APC tetramers: **(Fig. S8B-G)** cells from C57BL/6 mice at day 15 post-infection with influenza A/Puerto Rico/8/1934, from mice immunized with wild-type VSV at 10 days post-infection, and from mice immunized with rVSV/EBOV at 10 days post-infection.

### Quantification and Statistical Analysis

Significance was defined as p<0.05. Individual data points are shown. All statistics were calculated using GraphPad Prism 8 software (GraphPad Software, Inc). Statistical analysis was performed on log-transformed values for data represented on a log scale. All data shown on a log scale is represented as geometric mean ± geometric SD. Specific tests of statistical significance are detailed in figure captions and are as follows: For outcomes of ma-EBOV infection; statistical differences in survival data were analyzed by log-rank Mantel-Cox test, TCID_50_’s and body weight were analyzed via a two-way ANOVA with Tukey’s multiple comparisons test, and serum sGP was analyzed via a one-way ANOVA with Tukey’s multiple comparisons test. AUC measurements in all figures were analyzed via an unpaired student’s t-test. Statistical differences in quantities of anti-EBOV Ig shown via ELISA were analyzed by a two-way ANOVA or a one-way ANOVA with Tukey’s multiple comparisons test. Statistical differences in IgG binding avidity, IgG subtype measurements, neutralization assays, or ADCD were analyzed via two-way ANOVA with Tukey’s multiple comparisons test. Statistical differences in cell numbers acquired via flow cytometry were analyzed via an unpaired student’s t-test or a two-way ANOVA with Tukey’s multiple comparisons test. Statistical differences for quantified IF images were evaluated via an unpaired student’s t-test or a one-way ANOVA with Tukey’s multiple comparisons test.

## ACKNOWLEDGEMENTS

This study was supported by the National Institutes of Health (NIH, USA) grants R21AI139902 and UH2AI174415 to WM and NSB. NSB was also supported by NIH R01AI1125446 and R01AI167058. JA and TR were supported by NIH R01AI152476 and R01AI153413. JE was supported by NIH T32AI007485 and F30AI181340-01A1. AM and KLO were supported by an Intramural Research Program, NIH award (AI001254) to AM.

This research was supported in part by the Intramural Research Program of the National Institutes of Health (NIH). The contributions of the NIH authors were made as part of their official duties as NIH federal employees, are in compliance with agency policy requirements, and are considered Works of the United States Government. However, the findings and conclusions presented in this paper are those of the authors and do not necessarily reflect the views of the NIH or the U.S. Department of Health and Human Services.

We thank Jonathan Bernardi, Fionna Surette, and Peng Shao for their assistance in *Plasmodium* infections throughout the duration of the study. Special thanks to Hanora Van Ert, Paige Richards, and Caroline Plescia for their insight during weekly lab meetings. We also thank the University of Iowa Central Microscopy Research Facility (CMRF), a core resource supported by the University of Iowa Vice President for Research and the Carver College of Medicine.

## AUTHOR CONTRIBUTIONS

Conceptualization, JE, NSB, and WM; Methodology, JE, LG, KO, AM, NSB, and WM; Investigation, JE, LG, KO, CM, KR, JA, AT, RV, AM. Writing – Original Draft, JE; Writing – Review and Editing, JE, LG, KO, KR, JA, AT, AP, RV, TR, AM, NSB, WM; Funding Acquisition, NSB and WM; Resources, TR, AM, NSB and WM. Supervision; WM, NSB.

## FIGURE LEGENDS

**Figure S1: An acute, blood-stage *Plasmodium* infection decreases rVSV/EBOV-mediated protection against ma-EBOV challenge.**

Related to Figure 1.

Female C57BL/6 mice were infected with *Py* and vaccinated with rVSV/EBOV i.m. 6 days later. Mice were challenged 45-dpv with 10 FFU mouse-adapted Ebolavirus (ma-EBOV). Serum soluble glycoprotein (sGP) as evaluated by ELISA at 4-dpi (n=5/group, corresponds to experimental groups in Figure 1).

Data were analyzed with a one-way ANOVA and are represented as mean ± SD. p<0.05 considered significant.

**Figure S2: rVSV/EBOV-elicited antibodies are quantitatively reduced in experimentally *Plasmodium* infected vaccine recipients with minimal changes in antibody function.**

Related to Figure 2. Female C57BL/6 or Balb/c mice were *Py-*infected (red squares), *Pcc-* infected (green squares) or left non-infected (black circles) and 6 days later vaccinated as indicated in individual panels.

(A) BALB/c mice were vaccinated i.m. with 10^3^ PFU rVSV/EBOV and serum anti-EBOV GP IgG ELISA was performed at 21-dpv (n=5/group, data generated from one experiment).

(B) C57BL/6 mice were vaccinated i.m. with 6.25×10^7^ TCID_50_ rVSV/EBOV and serum anti-EBOV GP IgG ELISA was performed at 21- and 42-dpv. (n=5/group, data is representative of 3 independent experiments).

(C) C57BL/6 mice were vaccinated intraperitoneally (i.p.) with 2.5×10^5^ PFU rVSV/EBOV and serum anti-rVSV/EBOV Ig was assessed via indirect ELISA at 42-dpv. (n=10/group, data is pooled from 2 independent experiments and is represented relative to the highest serum concentration of anti-rVSV/EBOV Ig measured).

(D) anti-EBOV GP IgG and IgM were assessed at 42-dpv in 2.5×10^5^ PFU i.m. recipients via indirect ELISA (n=5/group, data is representative of 2 independent experiments).

(E) Indirect ELISA probing for anti-EBOV GP IgG from 42 days post vaccine of i.p. 2.5×10^5^ PFU rVSV/EBOV recipients in the presence of varying concentrations of ammonium thiocyanate (NH_4_SCN) to evaluate binding avidity. Data is represented as the percentage of total binding without NH_4_SCN treatment. (n=5/group, representative from 2 independent experiments).

(F) C57BL/6 mice were primed with 10^3^ PFU rVSV/EBOV delivered i.m., then boosted at 45-dpv with 2.5×10^5^ PFU rVSV/EBOV to elicit memory-derived antibody responses 5 days later (50-dpv). Serum anti-EBOV IgG at 45-dpv (before boost) and 50-dpv (after boost) are shown (left). Fold change in serum anti-EBOV IgG from before and after boosting is summarized (right).

(n=4-5 per group, data is representative of 2 independent experiments).

(G) C57BL/6 mice were primed with 2.5×10^5^ PFU rVSV/EBOV delivered i.p., then boosted at 42-dpv with 2.5×10^5^ PFU rVSV/EBOV to elicit memory-derived antibody responses 5 days later

(50-dpv). Serum anti-rVSV/EBOV Ig from before and after boost are shown (left). Fold change in serum anti-rVSV/EBOV Ig is summarized (right). (n=10/group, data generated from one experiment and is represented relative to the highest serum concentration of anti-rVSV/EBOV Ig measured).

Data in A and fold change in F were analyzed using a Student’s t-test and is represented as mean ± SD. Data in B-G was analyzed using a two-way ANOVA with Tukey’s multiple comparisons test and is represented as mean ± SD. p<0.05 considered significant.

**Figure S3: *Py* infection decreases lymph node germinal center responses to rVSV/EBOV.**

Related to Figure 3.

(A-B) Female C57BL/6 mice were vaccinated with 10^3^ PFU or 6.25×10^7^ PFU rVSV/EBOV i.m. and draining pLN GCBCs and GC Tfhs were assessed via flow cytometry at 0-, 6-, 8-, 10-, 14-, 18-, and 21-dpv.

(A) Representative flow plots for GCBCs are gated on CD19^+^B220^+^ single cells (left) and summarized cell numbers are shown per lymph node (right) (n=3/group, 1 independent experiment).

(B) Representative flow plots for GC Tfh cells are gated on CD90.2^+^CD4^+^CD44^+^ single cells (left) and summarized cell numbers are shown per lymph node (right) (n=3/group, 1 independent experiment).

(C, D) Female C57BL/6 mice were *Py* infected or left non-infected and vaccinated 6 days later with either 10^3^ PFU or 2.5×10^5^ PFU rVSV/EBOV i.m.

(C) GC Tfh cells assessed at 10-dpv. Representative flow plots are gated on CD90.2+CD4+CD44+ single cells (left) and summarized cell numbers are shown per lymph node (right) (n=7-10/group, data pooled from 3 independent experiments).

(D) Additional quantification of germinal center imaging from Figure 3C is shown: # of GCs per pLN (left); total GC area per pLN (middle); and total pLN area (right) (n=2-3/group, 3-6 sections per LN, data is pooled from 2 independent experiments).

Data in A and B are represented as mean ± SD. Data in C were analyzed using a Student’s t-test and is represented as mean ± SD. Data in D were analyzed using a two-way ANOVA with Tukey’s multiple comparisons test and is represented as the mean with bold dotted line, and light dotted lines representing SD. p<0.05 is considered significant.

Also see Figure S4.

**Figure S4: Gating strategies for flow cytometry of murine LNs and generation of EBOV GP tetramers.**

Related to Figures 3 and S3. Gating strategies are derived from non-infected IFN-γ^+/+^ vaccine recipients at 10-dpv.

(A-C) Gating strategy for identification of (A) poly-specific vaccine-elicited germinal center B cells (GCBCs), (B) poly-specific vaccine-elicited germinal center T follicular helper cells (GC Tfh), and (C) EBOV GP tetramer-positive GCBCs.

(D) SDS-PAGE analysis shows a major band corresponding to EBOV GP, confirming successful expression and purification. The lane marked ‘M’ contains molecular weight markers, with their approximate sizes indicated in kilodaltons. The lane labeled ‘EBOV GP’ shows the detected EBOV GP protein.

(E-G) Mediastinal lymph node cells from three groups of C57BL/6 mice were stained with EBOV GP-PE and EBOV GP-APC tetramers: (E) mice at day 15 post-infection with influenza A/Puerto Rico/8/1934, (F) mice vaccinated with wild-type VSV at day 10, and (G) mice vaccinated with rVSV/EBOV at day 10. Plots in show cells gated as live CD19^+^.

(H-J) Tetramer binding cells were identified in the IgM^-^GL7^+^ GCB cell population from (H) influenza A/Puerto Rico/8/1934 infected, (I) rVSV/G vaccinated and (J) rVSV/EBOV vaccinated.

**Figure S5: *Py* infection induces the expression of IFN-γ-associated genes in multiple cell types in the popliteal lymph node.**

Related to Figure 4.

(A, B) UMAP plot of B cells or T cells from each assayed sample in Figure 4 were extracted, re-clustered, and re-embedded in new UMAP coordinates. Cells are colored by assayed pLN sample; Day 0 *Py* infection (Naïve), teal; Day 6 *Py* infection, red; 24 Hour rIFN-γ-treated, blue.

(C, D) GSEA dot plot showing enriched gene sets in *Py*-infected versus naïve pLN B cells and T cells.

(E, F) Related to Figure 4G. List of log_2_fold >3 DEGs over naïve cells that are shared in pLN B cells or T cells from *Py*-infected and IFN-γ treated groups.

**Figure S6: *Py* infection causes an IFN-γ-dependent reduction in rVSV/EBOV GP-elicited antibody responses due in part to reduced vaccine replication.**

Related to Figure 5.

(A) Female IFN-γ^+/+^ or IFN-γ^-/-^ C57BL/6 mice were infected with *Py a*nd parastemia was evaluated from 8-25 dpi. Data are represented as a percentage of infected RBCs (iRBCs).

(B, C) C57BL/6 non-infected mice were vaccinated with 10^3^ PFU rVSV/EBOV and either 1×10^4^ TCID_50_ EBOVΔVP30 (B) or 10^3^ PFU UV-inactivated rVSV/EBOV (C). GCBCs were assessed at 10-dpv. Representative flow plots for GCBCs are gated on CD19+B220+ single cells (left) and summarized cell numbers are shown per lymph node (right) (n=1-7/group, 1 independent experiment).

(D) C57BL/6 mice were Py-infected or left non-infected and vaccinated 6 days later with either 10^3^ PFU rVSV/EBOV i.m. OR a high dose of EBOVΔVP30 sufficient to elicit comparable numbers of GCBCs in the draining LN at 10-dpv. Representative flow plots are gated on CD19+B220+ single cells (left) and summarized cell numbers are shown per lymph node (right) (n=4/group, data representative of 1 independent experiment).

(E) C57BL/6 mice were Py-infected or left non-infected and vaccinated 6 days later with either 10^3^ PFU rVSV/EBOV i.m. or a high dose (6.25×10^7^ PFU equivalent) of UV-inactivated rVSV/EBOV sufficient to elicit comparable numbers of GCBCs in the draining LN at 10-dpv. Representative flow plots are gated on CD19+B220+ single cells (left) and summarized cell numbers are shown per lymph node (right) (n=3-4/group, data representative of 1 independent experiment).

Data in A represented as the mean ± SD. Data in B and C were analyzed using a Student’s t-test and are represented as mean ± SD. Data in D and E were analyzed using a two-way ANOVA with Tukey’s multiple comparisons test and are represented as mean ± SD. p<0.05 considered significant.

Also see Figure S3.

**Figure S7: *Py* infection imposes an IFN-γ-dependent decrease in rVSV/EBOV replication in the draining LN.**

Related to Figure 6. Female C57BL/6 IFN-γ^+/+^ or IFN-γ^-/-^ mice were infected with *Py* and 6 days later vaccinated with either 10^3^ or 2.5×10^5^ PFU rVSV/EBOV i.m.

(A) C57BL/6 IFN-γ^+/+^ were injected i.m. with 10^6^ PFU rVSV/EBOV and draining pLNs were harvested at 12-hpv for immunoblot. Representative blot (left) and summarized optical density is shown (right) (n=1/group, data is pooled from 2 independent experiments).

(B) C57BL/6 mice were *Py-*infected or left non-infected and vaccinated with 2.5×10^5^ PFU rVSV/EBOV-GFP. Frozen sections of pLN at 6-hpv were imaged as shown (left) and % of GFP+ pLN area was quantified (right). Original magnification, x20. Green, GFP; blue, B220+ B cells. (n=2 per group, 2-4 sections per lymph node, data is pooled from 2 independent experiments).

(C) Quantification of frozen pLN sections harvested at 6 hours post vaccination with 2.5×10^5^ PFU from IFN-γ^+/+^ or IFN-γ^-/-^ mice from Figure 6F-G.

Data in B were analyzed using a Student’s t-test. Data in C were analyzed using a two-way ANOVA with uncorrected Fisher’s LSD test. Data in both B-C represent the mean with bold dotted line, with light dotted lines representing SD. p<0.05 considered significant.

